# Defining protein variant functions using high-complexity mutagenesis libraries and enhanced mutant detection software ASMv1.0

**DOI:** 10.1101/2021.06.16.448102

**Authors:** Xiaoping Yang, Andrew L. Hong, Ted Sharpe, Andrew O. Giacomelli, Robert E. Lintner, Douglas Alan, Thomas Green, Tikvah K. Hayes, Federica Piccioni, Briana Fritchman, Hinako Kawabe, Edith Sawyer, Luke Sprenkle, Benjamin P. Lee, Nicole S. Persky, Adam Brown, Heidi Greulich, Andrew J. Aguirre, Matthew Meyerson, William C. Hahn, Cory M. Johannessen, David E. Root

**Affiliations:** Genetic Perturbation Platform, Broad Institute of MIT and Harvard, Cambridge, MA 02142, USA; Cancer Program, Broad Institute of MIT and Harvard, Cambridge, MA 02142, USA; Data Science Platform, Broad Institute of MIT and Harvard, Cambridge, MA 02142, USA; Genomics Platform, Broad Institute of MIT and Harvard, Cambridge, MA 02142, USA; Department of Pediatrics, Emory University, Atlanta, GA 30322, USA; Aflac Center for Cancer and Blood Disorders, Children’s Healthcare of Atlanta, Atlanta, GA 30322, USA; Department of Medical Oncology, Dana-Farber Cancer Institute, Boston, MA 02215, USA; Department of Genetics, Harvard Medical School, Boston, MA 02115, USA

**Keywords:** saturation mutagenesis, deep mutational scanning (DMS), variants, open reading frame (ORF), log-fold change (LFC), base miscall, DNA oligonucleotide synthesis error, DNA polymerase error, next-generation sequencing (NGS)

## Abstract

Pooled variant expression libraries can test the phenotypes of thousands of variants of a gene in a single multiplexed experiment. In a library encoding all single-amino-acid substitutions of a protein, each variant differs from its reference only at a single codon-position located anywhere along the coding sequence. Consequently, accurately identifying these variants by sequencing is a major technical challenge. A popular but expensive brute-force approach is to divide the pool of variants into multiple smaller sub-libraries that each contains variants of a small region and that must each be constructed and screened individually, but that can then be PCR-amplified and fully sequenced with a single read to allow direct readout of variant abundance. Here we present an approach to screen very large variant libraries with mutations spanning a wide region in a single pool, including library design criteria and mutant-detection algorithms that permit reliable calling and counting of variants from large-scale sequencing data.

## Background

A pooled open reading frame (ORF) variant expression library for a gene of interest can be screened for cellular phenotypes, providing a powerful means to systematically characterize the functions of thousands of single-amino-acid substitutions in a single experiment. Advances in DNA oligonucleotide and gene synthesis [1] and in next-generation sequencing [2] have enabled production and screening of large variant pools. Pools encompassing all possible single-amino-acid changes for every amino acid position of a protein are known as saturation mutagenesis or deep mutational scanning (DMS) libraries. Screening such a library in cells can map each variant to phenotype in a single experiment.

As illustrated in **Figure 1**, a typical pooled screen tests the functions of thousands of genetic perturbation agents in a single experiment. In general, cell-based pooled screens rely on introduction of one genetic perturbation cassette, e.g., a CRISPR sgRNA or an ORF variant, per cell into the genome using delivery by a retrovirus, so that a single perturbation cassette is inserted into genomic DNA of each cell. The perturbed cell population is then subjected to selection pressure, for example, treatment with a drug, flow-sorting for reporter expression, or simply growth in culture. Post-selection cell samples are collected and processed to measure the abundance of library members in each sample by sequencing. Enrichment or depletion of each library perturbation in samples that have undergone selection versus samples under reference conditions (e.g., pre-selection or vehicle treatment samples) reveals the impact of the perturbations. For CRISPR screens, the functional elements themselves, the sgRNAs, serve as ideal short unambiguous barcodes to use for screen “deconvolution”, the measurement of enriched and depleted members of the library. The deconvolution of CRISPR libraries is a direct readout that simply counts sgRNA-matching sequencing reads. In contrast, ORF variant library screens are confronted with greater challenges for variant detection.

**Figure 1.**
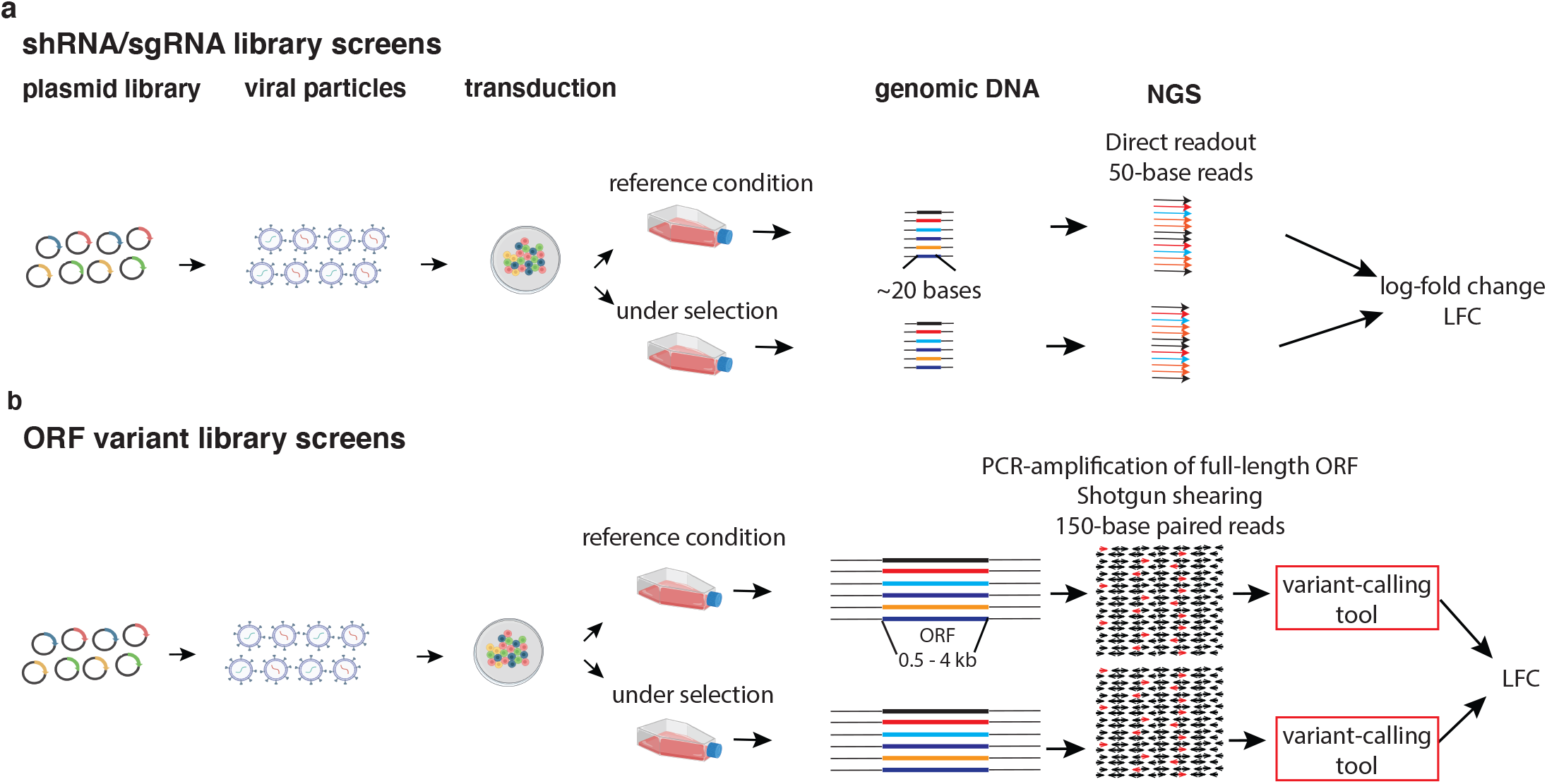
Pooled screens with ORF variant libraries versus sgRNA libraries. For either screen type, a plasmid pool of genetic perturbation agents is first packaged into lentiviral particles. Cells are transduced with the pooled virus at an infection rate of 30-50% to ensure that most of the infected cells get a single genetic perturbation. The perturbed cell population is divided into selection and reference arms. As the screen completes, the cell populations are harvested, and the collected screening samples are processed to quantify the relative abundance of the library perturbations for each sample, i.e., “deconvolution.” The relative enrichment or depletion of each perturbation in treatment versus reference condition is assessed differently for sgRNA versus ORF screens. **(a) A typical pooled screen using sgRNA libraries**. From the gDNA of collected cells, the guide sequence itself is PCR-amplified to serve as a barcode of the perturbation that was introduced to each cell. The sgRNA barcodes are short enough to be read straight out by a short NGS read. **(b) ORF variant library screens**. For an ORF variant library, the entire ORF is amplified from gDNA to serve as a ‘barcode’ of the variant. The distinguishing feature among barcodes is the mutation of a single codon position located anywhere along the entire ORF. The following steps serve to quantify the relative abundance of variants: (1) PCR-amplification to extract ORFs from genomic DNA, (2) random shearing of the full-length ORF, (3) next-generation sequencing of the resulting fragments, (4) application of variant detecting tools to identify and tally the reads. The NGS reads consist of variant signature-carrying reads (shown as red arrows) and wild-type reads (black arrows). The variant-calling software processes these NGS reads to determine the relative abundance of each variant and enables downstream analyses that map variants to phenotype strength, gauged by log-fold changes (LFCs).

The complexity of a site-saturated ORF variant library is a function of the ORF length and the number of desired variants per codon position. As illustrated in **Supplementary Figure 1**, at each amino acid position, the template amino acid is typically mutated to 19 other amino acids, a stop codon, and when possible synonymous codons that can serve as loss-of-function (LOF) and negative controls respectively (**Methods**). Factoring in the amino acid positions to be mutagenized (N), the library is a collection of approximately 20*N variants, or alleles, each differing from the template ORF at a single amino acid position. Capturing differences among the variant-defining signatures that are as subtle as a single codon variation located anywhere along the full-length ORF sequence is a significant technical challenge. A popular approach has been to employ multiple sub-libraries, each with short mutated regions with lengths of 25 [3, 4], then 47 [5], 100 [6], and recently as many as 170 [7] amino acids. These sub-libraries were constructed, screened and sequenced separately. Each of these sub-libraries covers a mutated region short enough to be covered by a single next-generation sequencing read for direct readout of variants. While practical for small ORFs, for longer genes this approach requires an unwieldy number of mini-libraries and screens. Our methodology, on the other hand, permits screening of very large pools in a single experiment using robust variant-calling tools that can accurately quantify variants from sequencing data generated by amplifying and randomly shearing the entire ORF. A single site-saturated variant library is thus adequate a wide range of ORF lengths, e.g., ~600-bp *KRAS* [8], ~1200-bp *TP53* [9] and *SMARCB1*, ~1400-bp catalytic region of *PDE3A* [10], ~1700-bp *SHOC2* [11], ~1800-bp *PPM1D* [12], ~2200-bp *EZH2*, ~3600-bp *EGFR*) (**Fig. 1b**). Here, we present methods and strategies developed in the process designing, constructing and screening more than 20 variant libraries (**Supplementary Table 1**). We have made available our library designer, variant-calling software, and data analysis tools (**Supplementary Table 2)**. We anticipate that these tools and methods will facilitate variant screening for longer ORFs.

## Results

### Variant calling with ASMv1.0

The challenge of calling variants correctly from PCR-amplified, fragmented, and sequenced genomic DNA stems from multiple factors that lead to unexpected or misattributed variant sequences. In addition to the intended, library-designed codon sequence modifications relative to the reference sequence, each ORF molecule in our library is subjected to two kinds of unintended changes that occur in the process of library construction: oligonucleotide chemical synthesis errors and DNA polymerase errors. Although the detailed synthesis strategies can vary, ORF variant libraries are typically all produced using chemically synthesized oligonucleotide strands that encode the variant-defining codon modifications and that constitute part of the final full ORF library sequences, by coupling with unmodified reference (often “wild-type”) portions of the ORF that are enzymatically synthesized by high-fidelity DNA polymerases in a PCR process. To fully characterize the composition of a variant library, it is desirable to identify empirically all the observed sequence variations between the reads and the reference sequence, without consideration of the intended library design. In general, these variations will include nucleotide substitutions, insertions, deletions and combinations thereof. Some variations exactly match design-intended variants, while others carry unexpected nucleotide changes in addition to, or instead of, the expected variation. Aside from the initial library production process, the process of extracting and sequencing ORF molecules from the library pDNA or the genomic DNA of screened cells, a process referred to as “library deconvolution”, is itself subject to errors from imperfect PCR amplification, and from sequencing miscalls (e.g., a Phred Q20 base-call has a 99% chance of being correct). Our enhanced variant-detection software aimed to distinguish the intended variants from those arising from errors of synthesis of the initial library or from the library “deconvolution” process, and to assess library quality and deconvolution miscall rates.

Improving upon our earlier version of variant-calling strategy and the associated software (ORFCallv1.0), we developed Analyze Saturation Mutagenesis v1.0 (ASMv1.0; see **Methods** and **Supplementary Fig. 2**) to more accurately interpret pooled variant NGS data and measure variant abundance. ASMv1.0 is now included in the Genome Analysis Toolkit (GATK v4.2.0.0). NGS reads are each first aligned to the reference (or template) ORF to produce binary alignment map (BAM) files that constitute the input to ASMv1.0. ASMv1.0 then performs all the steps required, starting from per-sample read-and-reference alignments, to produce a list of observed nucleotide sequence variations in relation to the reference ORF and the abundance of each of these nucleotide sequence variations for each sample. Notably, the list of observed nucleotide sequence variations is independent of any prior expectations of the library’s variant composition. As shown in **Supplementary Figure 2**, the key steps in this process are: (1) trimming from the reads the flanking sequences introduced during fragmentation and barcoding of the ORF DNA, (2) combining overlapping paired-end reads when possible to obtain a “per-molecule” single sequence result per read pair, (3) applying base quality, minimum read length and other exclusion filters prior to variant calling, (4) identifying bona fide sequence variations, if any, for each read or read pair versus the reference (substitutions, insertions, deletions, or a set thereof), and (5) counting examples of each set of observed nucleotide sequence variations (note in this report, a variant is defined by a set of nucleotide sequence variations), to quantify the abundance of that variant in each screening sample. Because the NGS reads are far shorter than the full-length ORF and because each planned variant in the library typically carries only a single codon change from the template ORF, most observed aligned reads will exactly match the reference sequence. These perfectly reference-matching reads are used to assess the depth of read coverage across the reference ORF and this read coverage is used to normalize the variant abundance. The reads that contain mutations provide the identity and abundance of each variant sequence that is defined by those mutations. Because the ASMv1.0 algorithm identifies empirically observed variants in an entirely data-driven manner that does not rely on expectations based on the library design, the output can be used not only to filter for the results for designed ORF variants (i.e., the intended variants), but also to assess the library quality and characterize its actual composition, which will always differ to some extent from the intended design.

### The enhanced performance of ASMv1.0

ASMv1.0 incorporates several enhancements versus an earlier variant-detection process, implemented in the software package ORFCallv1.0. ORFCallv1.0 is a simple and effective variant detection tool designed strictly for single amino acid substitution variant libraries, and it was employed to interpret a number of saturation mutagenesis screen results [9, 13–15]. ORFCallv1.0 quantifies variant abundance by first aligning reads to the reference sequence, then recording at each codon position the count of each of the 64 possible codons. Note that ORFCallv1.0 tallies the codon read counts at each amino acid position by staying within the 3-nucleotide codon window and ignoring any additional changes located in the same read. If a library has high rate of synthesis errors (e.g., due to oligonucleotide chemical synthesis errors), these errors will lead to high frequency of unintended changes that are combined, in the same molecule, with an intended codon change. Ignoring the presence of these unintended changes in the manner of the ORFCallv1.0 algorithm can be costly when these additional nucleotide changes produce phenotypes that differ from the phenotype of the pristine intended variant. Because ASMv1.0 calls variants in the context of full-length read pairs, incorporating all nucleotide variations - substitutions, insertions and deletions - observed between the read pair and the reference sequence, it can separate pure variants that bear only the intended sequence changes from those that also harbor extra changes. **Supplementary Table 3** summarizes the enhanced functionalities from ORFCallv1.0 to ASMv1.0.

We used the screens performed in Giacomelli *et al* [9] to compare the two variant-detection methods. Ectopically expressed *TP53* variants were screened in isogenic p53^WT^ and p53^NULL^ A549 lung cancer cells, with selection pressure from either nutlin-3 or etoposide. Under nutlin-3 treatment, the screen of p53^NULL^ cells enriched *TP53* loss-of-function (LOF) variants and depleted wild-type-like variants (**Fig. 2**). Under etoposide treatment, the p53^NULL^ cells enriched wild-type-like *TP53* variants and depleted the *TP53* LOF variants (**Supplementary Fig. 3**). Under nutlin-3 treatment, the screen of p53^WT^ cells enriched *TP53* variants exhibiting dominant-negative effects (DNE) (**Supplementary Fig. 4**). Using both the ORFCallv1.0 and ASMv1.0 variant-calling methods, we reprocessed the next-generation sequencing data from these three screens. The phenotypic strength of each variant, measured as log-fold change (LFC) between pre- and post-treatment samples, produced similar heatmaps, indicating that the two variant-calling algorithms are very consistent with each other for most variants (**Fig. 2a, Supplementary Figs. 3a, 4a**). We noted however that a small subset of variants (~3% of the library) exhibited substantial differences between the LFC values calculated using these two algorithms (delta(LFC)=LFC^ASM^-LFC^ORFCall^, **Fig. 2b, Supplementary Figs. 3b** and **4b**).

**Figure 2.**
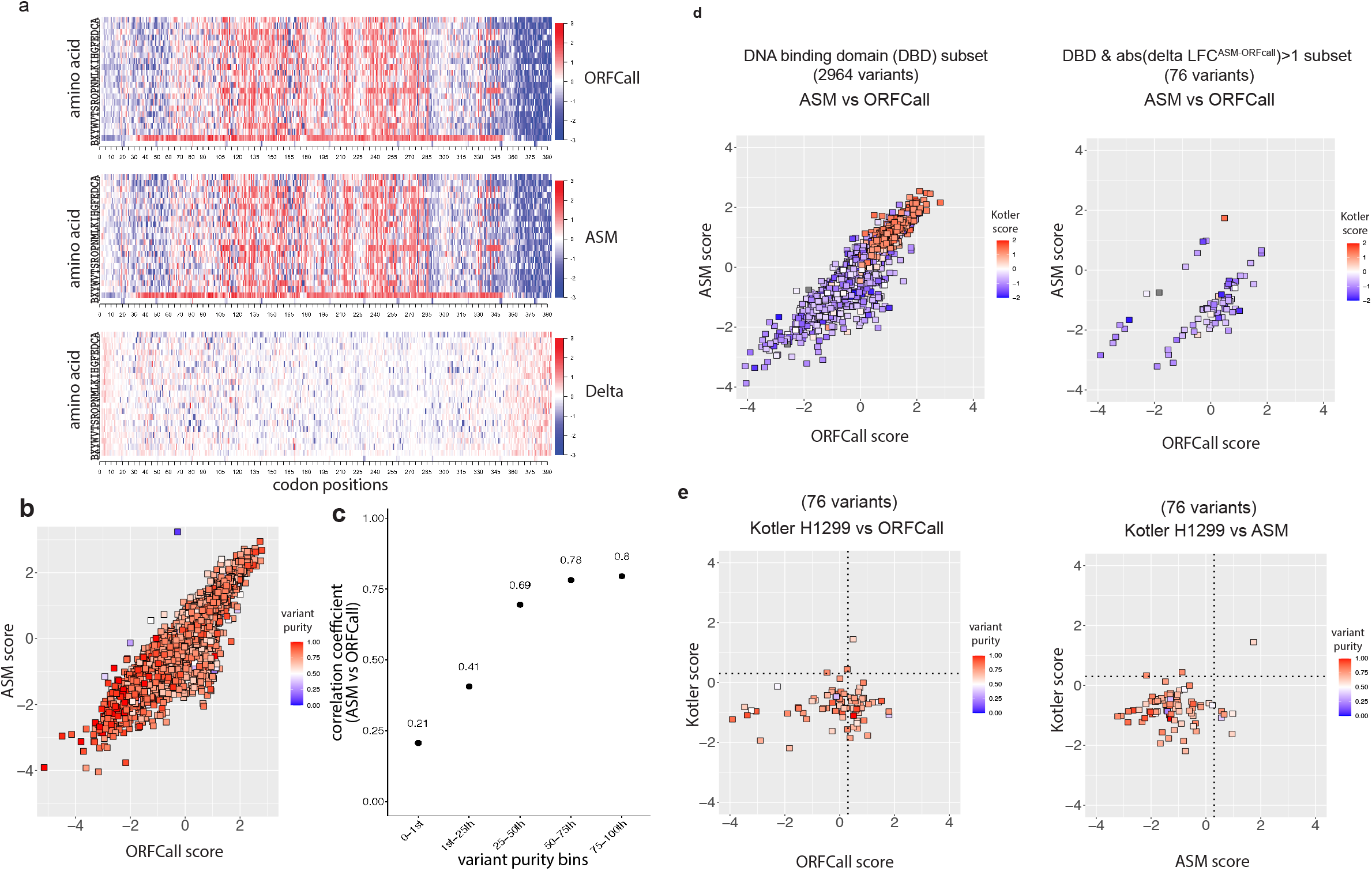
Reanalysis of *TP53* saturation mutagenesis screen using ASMv1.0. The Giacomelli *et al* [9] p53^NULL^ A549 cell/nutlin-3 screen enriches *TP53* variants that exhibit loss-of-function (LOF), and depletes wild-type-like variants. (**a) LFC heatmaps depicting enrichment or depletion of all variants in this library**. Each variant is placed in a 2-dimensional grid, where the horizontal dimension specifies amino acid position, and the vertical dimension specifies the amino acid substitution. ‘X’ designates nonsense mutations, and ‘B’ designates silent codon changes. The heat index is log-fold change (LFC) between treatment and reference samples. The same original NGS screen data were analyzed using ORFCallv1.0 (top) and ASMv1.0 (middle), yielding mostly similar results. The small subset of variants for which the two analyses give different results are highlighted in the third heatmap (bottom) that shows delta(LFC)=(LFC^ASM^ – LFC^ORFCall^) as the heat index. (**b**) **Scatter plot comparing LFC values of all variants as assessed by the two versions of software**. The color index of each data point is the purity of the corresponding variant. (**c**) The percentile of “Variant purity”, calculated by ASMv1.0, was used to group variants into five bins. For each bin of variants the Pearson correlation coefficient between the ORFCallv1.0 and ASMv1.0-computed LFCs is plotted on the y-axis. (**d,e**) **Testing ASMv1.0 using the published data by Kotler *et al* [5]**. Despite the noted differences, the Kotler screen of p53^NULL^H1299 cells with a library of *TP53* DNA binding domain mutants provides an intriguing comparison to the larger Giacomelli pooled screen. For the 2964 variants in the Giacomelli library that are shared with the smaller Kotler library, 3-way comparisons are shown, with ORFCall and ASM scores as x- and y-axis, Kotler score as the color index (**d**, **left**). The 76 variants that overlap between the two studies and that scored most differently in the Giacomelli screen processed by the two versions of software are isolated **(d, right**). the Kotler scores compared to the corresponding Giacomelli LFC scores called by ORFCallv1.0 (**e, left**) and ASMv1.0 (**e, right**).

In a perfect library consisting solely of intended variants, ORFCallv1.0 and ASMv1.0 should deliver the same result. However, owing to errors introduced during library synthesis such as oligonucleotide synthesis errors and DNA polymerase errors incurred in the production of the ORF variant materials, it is inevitable that planned codon changes are sometimes physically accompanied by one or more unintended accidental nucleotide changes in the same molecule. We define the ‘variant purity’ of a designed variant as the ratio of molecules that exhibit exclusively the intended mutation to the sum of all molecules that harbor the intended codon change, regardless of whether they also harbor additional nucleotide changes. When the purity of a planned variant is high, both versions of the software will call and quantify variants similarly; however, when the variant purity is low the enrichment and depletion scores measured by ORFCallv1.0 can be misleading. We computed the purity of every planned variant using the reference samples (samples without selection pressure) from the Giacomelli screens, and for each purity bin we computed the Pearson correlation coefficient of two sets of LFCs, one using ORFCallv1.0, the other using ASMv1.0 (**Fig. 2c, Supplementary Figs. 3, 4)**. As expected, the two methods correlate well when the variant purity is high and diverge as the variant purity decreases.

Work by Kotler *et al* [5] offered an additional way of testing the performance of ASMv1.0 and ORFCallv1.0. We compared the Giacomelli screen LFC scores called by both versions of the software with the results from the Kotler *et al* study [5], which screened p53^NULL^ H1299 cells with a smaller variant library that was focused on the p53 DNA binding domain (DBD). Kotler *et al* targeted the p53 DBD with four sub-libraries, each covering a 47-amino acid region of the ORF. The four sub-libraries were screened separately, and the screen samples were processed by direct sequencing readout of the mutated region in each of the 4 libraries. Like the Giacomelli p53^NULL^ A549/nutlin-3 screen, the Kotler p53^NULL^ H1299 cell screen enriched for *TP53* alleles that exhibit LOF and depleted wild-type-like variants. From the 7890-variant Giacomelli p53^NULL^ A549/nutlin-3 screen data, we extracted a 2964-variant subset that covered the same amino acid mutational space as the four combined Kotler H1299 screens. These 2964 Kotler scores and the Giacomelli scores, using either ASMv1.0 or ORFCallv1.0, exhibited good correlation (Pearson coefficient 0.65 (p-value=0), and 0.62 (p-value=2.4e-317) respectively), despite the fact that the screens were performed using different variant detection methods, different libraries, different cell lines, and different conditions consisting in one case of simple growth in culture and in the other nutlin-3 treatment (**Supplementary Fig. 5**). Interestingly, when focusing on p53 DNA binding domain, both the Kotler and Giacomelli data exhibit a bimodal pattern representing the enriched p53 loss-of-function variants (**Supplementary Fig. 5, left,** upper right quadrant) and the depleted p53 wild-type-like variants (**Supplementary Fig. 5, left,** lower left quadrant). There is therefore a natural cutoff to classify each of the 2964 variants as having either LOF or WT-like activity. **Supplementary Figure 5** suggests that we may use score 0.3 as the cutoff for classification. Consistent with the high similarity noted above between the ASMv1.0 and ORFCallv1.0 scores for most variants, the agreement between Kotler and Giacomelli classifications is similar whether using the ASMv1.0 method (2519 out of 2964 variants, 85% agreement) or the ORFCallv1.0 method (2482 out of 2964 variants, 84% agreement). This is to be expected for a library that contains mostly high purity variants for which ASMv1.0 and ORFCallv1.0 show high agreement. However, focusing on the small subset of 76 variants that were scored very differently by ASMv1.0 and ORFCallv1.0 in the Giacomelli data (abs(delta(LFC)) >1, **Fig. 2d, right**), the ASMv1.0 classification agrees with the Kotler classification much better (69 out of 76 variants, 91%) than the ORFCallv1.0 classification does (42 out of 76 variants, 55%, **Fig. 2e**).

### ASMv1.0-enabled analysis reveals that the secondary mutations (errors) in the library are near the targeted mutation and are introduced by errors in DNA oligonucleotide synthesis

The ability of ASMv1.0 to present molecules with a full list of observed changes that include intended and/or unexpected changes along the entire read pair allows in-depth analyses of ORF variant library composition. Thus, in addition to the variant purity discussed earlier, we can also tally and characterize the unexpected nucleotide changes to profile the errors introduced during library construction. For example, we were able to determine that in the *TP53* library, 84% of molecules are pure and planned variants (**Fig. 3a, top**), and the errors in the library are predominantly single nucleotide changes.

**Figure 3.**
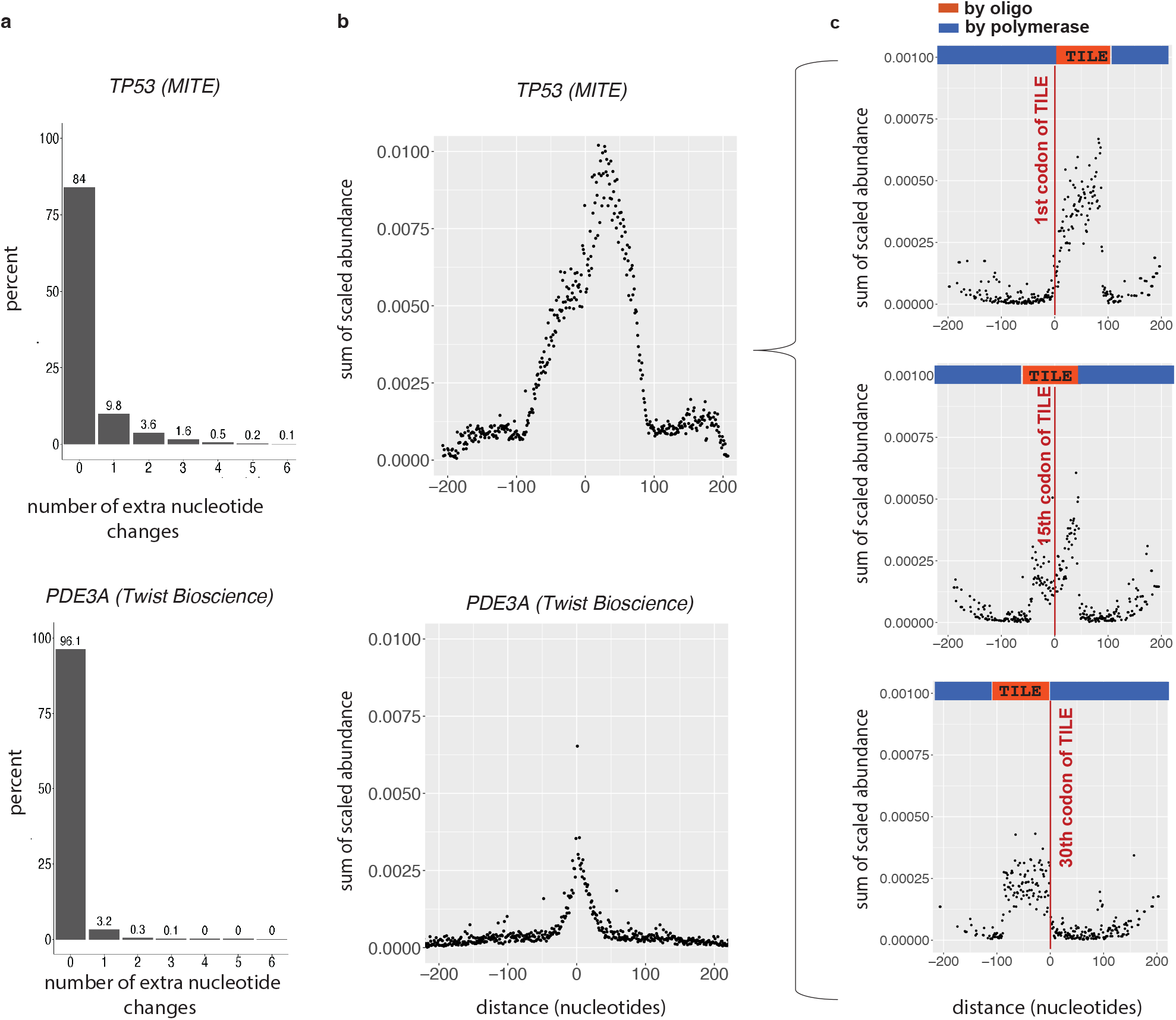
Unexpected mutations in the site-saturation variant libraries are rare and shown to co-localize with planned mutations. (**a**) **Breakdown of the planned variants versus those with additional mutations**. ASMv1.0 output records information that allows us to identify the number of additional unintended nucleotide changes that are associated with a planned codon change. The number of additional unexpected nucleotide changes were counted and the frequencies of different numbers of additional changes were shown in the bar-chart. The ‘0’ extra nucleotide changes category represents the pure variants. (**b**) **Unexpected mutations are concentrated in the vicinity of a planned change**. The ASMv1.0 output enables a tally of unexpected nucleotide changes as a function of their distance, in nucleotides, from the planned codon change. For all called variants that contained a planned codon change **and** additional changes (therefore these variants are all unexpected), the counts of these unexpected variants were first scaled to the fraction of the reference read coverage across the sequence variations that define the variants; scaling removed fragmentation bias as part of Illumina Nextera ORF sequencing process. These unexpected variants were then grouped by the distance between the error and the intended codon change, and the relative abundances in each distance group were summed for the group abundance total. Two libraries made with different technologies, *TP53* library by MITE and *PDE3A* by Twist Bioscience were compared. The MITE technology uses synthesized DNA strands of 150 bases long, of which 90 bases (called TILE) go into the final products, while Twist Bioscience introduced desired codon changes through oligonucleotide primers with the intended codon changes placed in the middle and flanked with 15-25 bases on either side. The major errors are concentrated within ~100 bases (MITE method) or ~30 bases (Twist Bioscience method) from a planned codon change. (**c**) The *TP53* library offers a clear boundary between chemically synthesized and polymerase copied nucleotides. The dataset used in **b** (above) was filtered in such a way that the intended codon modifications presented in these unexpected variants were those whose targeted codon positions placed them at the beginning, middle or end of a tile. Each of the three subsets was profiled for error-versus-position and each had clean crossings between chemically synthesized (region colored in red) and polymerase copied nucleotides (region colored in blue). We observed that errors are present predominantly in nucleotides synthesized chemically and the region produced by DNA polymerase can be treated as error-free.

Using ASMv1.0 output, we also investigated the proximity distribution of errors in relation to the intended codon change. In **Figure 3**, we compared libraries made by two different methods: a *TP53* library made by MITE (Mutagenesis by Integrated TilEs) [15], and a *PDE3A* library synthesized by Twist Bioscience [10]. The MITE method used chemically synthesized oligonucleotide strands to encode 90-nucleotide variable ‘tiles’ that were subsequently incorporated with the rest of the reference ORF segments to form the full-length ORF sequence. These reference segments were prepared by PCR with a high-fidelity DNA polymerase. The Twist Bioscience technology involved the chemical synthesis of oligonucleotides varying in length between 30-50 nucleotides, depending on GC content, with the variant-encoding bases located in the middle of the oligonucleotide; these oligonucleotides were used as the PCR primers and became part of the ORF sequences of the library. We profiled the library errors (i.e., unexpected nucleotide changes) as a function of the number of bases between the locations of the error and the intended codon change. Among variants that carry an intended codon change **and** one or more extra nucleotide errors, we grouped these variants by the distance between the location of the error and the closer edge of the intended codon. For each distance group, the scaled variant abundances (i.e., a variant count is scaled by dividing it with the read coverage across the region spanning the variant signature. See **Scaling** in **Methods**) were summed as the group abundance (**Fig. 3b**). We observed that the stretches of frequent errors in libraries produced by the two methods, ~90 bases for the MITE method and ~30 bases for the Twist Bioscience method, correspond faithfully to the length of chemically synthesized DNA oligonucleotides involved in library production. Since most of the errors in an ORF variant are concentrated near the intended codon change, sequencing reads as short as 100 bases can detect most of the unexpected changes that accompany each intended change.

It is important to note that all nucleotides in variant ORF libraries come from either of two sources: (1) chemically synthesized oligonucleotides that encode and introduce the mutations (e.g., PCR primers or oligonucleotide tiles that encode variants), or (2) nucleotides copied from the reference template by high-fidelity DNA polymerase. The errors in these libraries are therefore expected to arise predominantly from oligonucleotide chemical synthesis errors (~2 errors per kb), since DNA polymerase (Q5, Fushion, or Ex Taq) nucleotide misincorporation rates are far lower [16]. We demonstrate in **Figure 3c,** that this is exactly the case: The tile-nature of *TP53* library offers clear boundaries between chemically synthesized and polymerase incorporated nucleotides. We extended the error-versus-distance profiling by filtering the data used in **Figure 3b** to obtain three subsets, each respectively carrying intended codon changes targeting codon positions such that in the library production scheme these intended codon modifications were placed at the start (1^st^), middle (15^th^) and end (30^th^) of the 30-codon tiles. The three error-and-distance profiles all indicate that the errors in the libraries are confined in chemically synthesized oligonucleotides and as across the boundary into polymerase copied nucleotides the rate of errors drop precipitously. Altogether, next-generation sequencing reads of 100 bases or longer that carry the expected mutations, which mark those reads as those covering chemically synthesized nucleotides, will effectively also capture most of the accompanying synthesis errors in the library.

### ASMv1.0-enabled analysis assesses the effect of miscalls (artifactual errors) introduced by processes involving PCR

The synthesis errors discussed above are incorporated into the variant libraries and the resulting variants, although not part of the intended library design, are nonetheless actually present in the library and are screened in the cell populations to which the libraries are applied. It is important also to consider errors in the variant quantification process itself, i.e., miscalls that do not accurately reflect the variants present in the library. To deconvolute the variant abundances from a pooled library, PCR was used to replicate the ORF DNA from the genomic DNA of the library-transduced cells (up to 30 PCR cycles) and was used again in to barcode samples in preparation for NGS (12 cycles). We define ‘miscalls’ as the artifactual nucleotide calls that do not accurately reflect a sequence that was actually present in the sample. In this work, miscall rates are driven by the error rate of the DNA polymerases used for PCR to extract ORF sequence from gDNA and in the subsequent sequencing sample preparation, and also the sequencing error rates which are reported as Phred Q scores. To directly assess the rate of miscalls in the deconvolution of a variant library, we processed and sequenced a pure clonal reference *TP53* ORF sample that should therefore produce only reads that perfectly match the reference sequence. We observed miscalls at a rate of ~3-6 miscalls per 10,000 sequenced bases (**Fig. 4a, right**).

**Figure 4.**
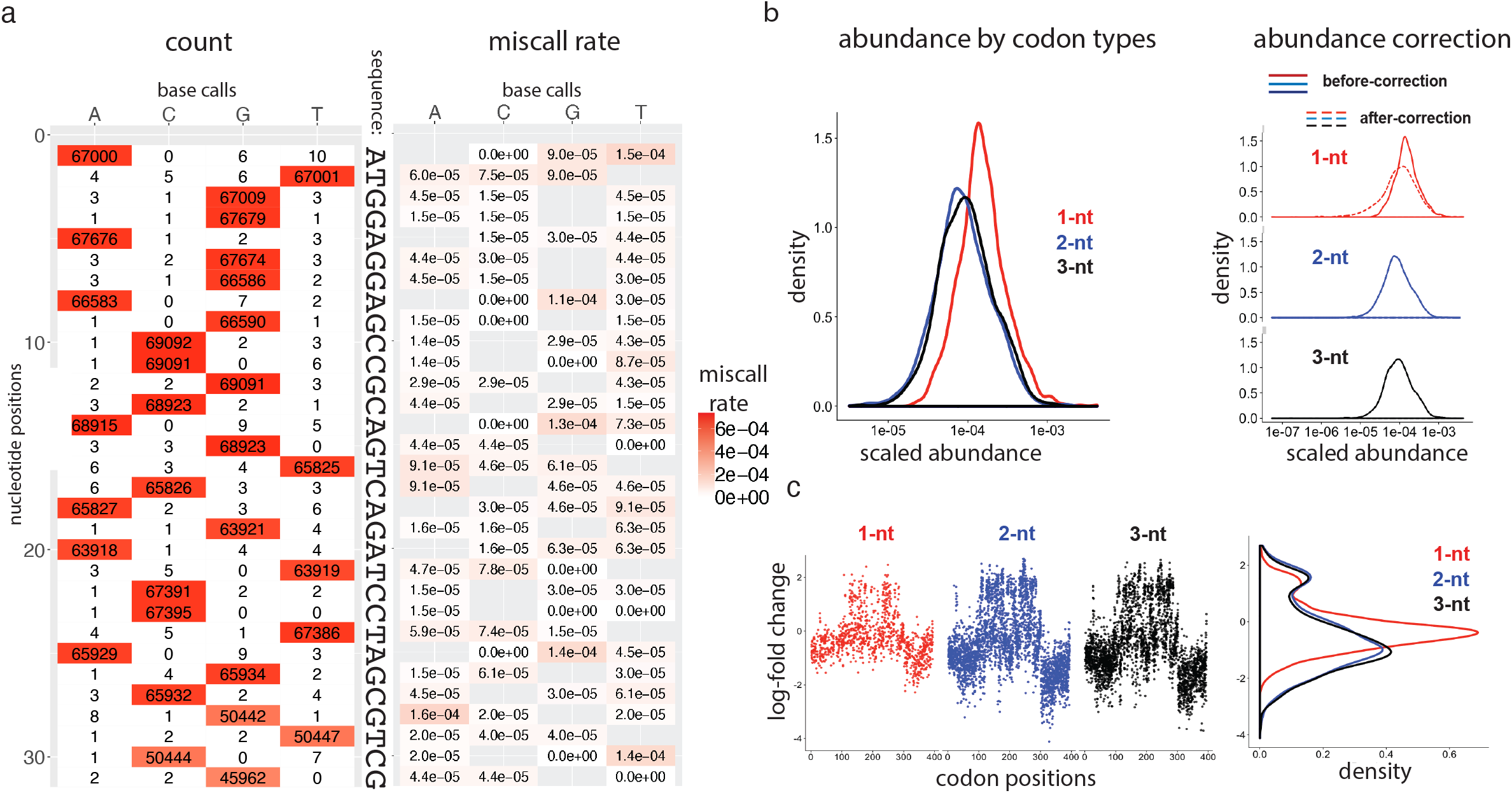
Miscalls due to PCR and sequencing errors and their effects on variant quantification and LFC. **(a)** A clonal *TP53* reference plasmid (i.e., all sequences are wild-type) was processed as if it were a screening sample, subjected to PCR amplification, NGS sample preparation and NGS sequencing read out. As expected, the base call at each position is predominantly, but not 100% uniformly, the reference nucleotide. The non-reference calls represent miscalled variants that were not actually present in the sample being measured. The heatmap on the left shows tallied counts of called nucleotides at each nucleotide position. The heatmap on the right depicts the rate of miscalls; blank spots in the heatmap are the wild-type nucleotides. (**b**) **Abundance distributions of 3 codon types in a reference screening sample**. *TP53* variants were grouped by the number of differing nucleotides between the variant signature codon and the reference codon; the 3 groups are called 1-nt-delta, 2-nt-delta and 3-nt-delta variants. The miscalls resulting from PCR/NGS errors inflate the abundance measurements. In the *TP53* library screen data, predominately affected are variants whose signature codon is 1-nt (but not 2- or 3-nt) away from the reference codon. As a result, the variants of 1-nt delta codons as a group appear more abundant than groups of 2- or 3-nt delta codons (**left**). The artifactual ‘variant’ abundance resulting from PCR/NGS errors can be corrected by subtracting the miscall rate as determined by a control experiment using a clonal sample from every screening sample (**right**). The variant abundance measurements of the 2-nt and 3-nt delta codons are much less affected by the miscalls and therefore the corrections do not appreciably alter the abundance distributions of these 2 codon types. (**c**) **Suppression of fold-changes in 1-nt delta codon variants**. Three scatter plots, one for each of the three codon types, depict LFC versus variants ordered by codon positions. A density plot on the right depicts the LFC score distribution within each of the three codon types. Using the pMT025-*TP53* library as an example, 1-nt delta variants (data series colored in red) showed narrower fold-change values than 2-nt and 3-nt delta variants.

The miscall rate is sufficiently large to be a significant factor in variant detection for large complex libraries. As an example, illustrated in **Supplementary Figure 6**, miscalls lead to (1) inflated counts of variants with codons that are one-nucleotide different from the template codon (we refer to this variant type as ‘1-nt delta’ codon), and (2) as a result of these inflated counts, artifactually reduced log-fold changes between samples of 1-nt delta codons that are truly present in the library (**Supplementary Fig. 6,** legend). Indeed, in the *TP53* library screen with p53^NULL^ cells/nutlin-3, we observed the inflation of counts (**Fig. 4b**), and the suppression in fold-change (**Fig. 4c**), in the 1-nt delta variant group relative to 2- or 3-nt delta variants. This contribution of PCR errors to 1-nt delta codon change counts has been observed in every library we have processed (~20 to date), and the effect increases as ORF length increases (data not shown).

The effects of PCR/NGS miscalls on the quantification of variants reshaped our strategy for library design rules and led us to add control experiments. First, we avoid the use of codons that are one nucleotide away from the reference codons whenever possible. For a typical ORF sequence, to achieve all possible single site amino acid substitutions, it is sometimes necessary to use codons that deviate from the corresponding reference codons by one nucleotide. As result of this design rule that limits the use of 1-nt delta codons, variants of 1-nt delta codons can be reduced to approximately 3% (from otherwise ~15%) of the library members. Secondly, to better interpret the results of those 3% remaining 1-nt delta variants, we perform the deconvolution process on a control sample that is a single clone of the reference ORF and therefore contains only one uniform sequence. This sample is processed exactly as if it were a screening sample. These clonal reference ORF data may be used to identify the 1-nt delta variants that are most affected by miscalls, or can even be used to apply a correction to the screen sample data based on the background miscall rates determined from this sample (**Fig. 4b**, **right**).

### Silent variants are used as a measure of screen baseline

The inclusion of silent variants provides a set of controls that serve as independently measured pseudo-replicates of the reference amino acid sequence. Like the missense- and nonsense-variant sequences, in order to reduce contamination by miscalls, these silent mutations should employ codons that are 2- or 3-nt different from the reference codon. Often, codons encoding the same amino acid differ from one another at the wobble base. As a result, many of the possible silent variant codons are one nucleotide away from the template codon but these are less useful as controls. Silent variants with 2- or 3-nt delta differences relative to their respective reference codon results in silent variants comprising about 2% of the library. As demonstrated by the bimodal split of phenotypes in *TP53* DBD library screens discussed earlier, ORF variant library screens often produce a high rate of “hit” variants that deviate from the reference-sequence phenotype. As a result, in contrast to many other common pooled screening situations, one cannot always rely on the large portion of the library to match the reference phenotype and to serve as a reference baseline, making the silent codon change controls a particularly important reference point for interpreting the screen results. The silent-variant LFC results are displayed in the row marked by ‘**B**’ in the heatmaps in **Figure 2a**.

In the *SMARCB1* library, we included nearly all possible silent variants, including 1-nt delta silent variants. This case provides an example of the consequences of single-nucleotide miscalls on the reference sequence that masquerade as 1-nt library variants. As expected, due to the miscalls discussed earlier, the apparent abundance of the library’s 1-nt delta silent variants is inflated by those miscalls relative to the abundance of 2- and 3-nt delta silent variants (**Supplementary Fig. 7a**). As a consequence, relative to 2- and 3-nt delta groups, the 1-nt delta silent variants show an artificially suppressed log-fold change range when comparing among samples, because every sample across which log-fold change comparisons are made receives the same contribution of counts due to miscalls of the reference sequence (**Supplementary Fig. 7b**), inflating and equalizing the counts for these 1-nt variants across samples. Our current downloadable library designer (**Supplementary Table 2**) avoids 1-nt delta silent variants and minimizes selection of 1-nt delta missense and nonsense variants.

### Using next-generation sequencing (NGS) versus long-read sequencing (LRS) for screen deconvolution

Long-read sequencing technologies have the potential to capture all mutations present on each ORF-spanning DNA molecule. However, the technology that can capture the readout of full-length ORF sequences [17, 18] has significant limitations for variant detection due to low concordance rate of base calls [19]. For example, in a study by Wenger *et al* [20], circular consensus sequencing (CCS), the best of its kind, has a concordance rate of 99.772%. This is equivalent to a Q26 Phred score, producing one miscall per 439-base readout. This sequencing-error rate is higher than the error rate in commercial DNA oligonucleotide synthesis, the key contributor of the real errors in the libraries as discussed earlier. As a result, one may be forced to either throw out the many reads that will show apparent secondary mutations due to miscalls or to focus locally on the expected codon changes and ignore surrounding mutations.

Long-read sequencing technology is advancing at a steady and rapid pace, and its applications are actively being explored. For example, based on circular consensus sequencing (CCS), by concatenating multiple DNA molecules, MAS-seq can increase the CCS throughput by 15 to 30-fold while maintaining the sequencing phredQ [21]. In the recent work by Frank *et al* [22], long read sequencing was used to associate a set of unique external barcodes to each of ~7000 variants in a saturation variant library, so that the screen deconvolution work can be achieved by direct next-generation sequencing of the barcodes. However, there are several factors that create a high bar for long-read sequencing to match the results obtained with NGS paired-end reads of 150 bases each with a Phred score >Q30, which is comfortably achievable by most NGS platforms: (1) We have demonstrated that the errors in our libraries are near the intended codon changes (**Fig. 3b**), and a 150-base read is adequate to detect both the variant-defining sequences and library errors from DNA oligonucleotide synthesis. We have shown that it is unnecessary to sequence an ORF variant into the regions synthesized by high-fidelity DNA polymerase during library construction, because the errors introduced by DNA polymerase in regions outside of a 100-base window around the intended codon changes are rare enough to be inconsequential (**Fig. 3c**). ASMv1.0 output offers many choices to filter the results in various ways, such as to retain only the reads that bear a library-designed codon change and that effectively represents the variant’s error-bearing region, i.e., the region that was chemically synthesized. (2) We have demonstrated that PCR-introduced miscalls produce predominantly 1-nt delta codon changes. Because we minimize the use of 1-nt delta codon variants, the artifactual variants resulting from miscalls are mostly unexpected variants that were not part of the library design. In this case, ignoring reads carrying only unexpected variations largely removes the effects of sequencing miscalls. (3) Even with high base-quality threshold, NGS currently produces adequate read yield for sequencing screen samples of variant libraries. ASMv1.0 allows users to set adequate base quality threshold, for example, at Phred score Q37 with Illumina Novaseq NGS. With this filter, we can comfortably achieve <1 miscall per 5000-base NGS readout. At this low miscall rate, >94% (0.9998^300) of 150-base read pairs used to call variants are free of miscalls.

## Discussion

Variant detection software ASMv1.0 implements in-depth data analyses that allows exploration of variant library composition and the fidelity of the screen sample deconvolution process. The types and occurrence frequency of unintended species that arise during the variant library production process and the overall variant purity of each designed library variant were empirically assessed for multiple ORF libraries. Miscalls arising from PCR and sequencing in the sample deconvolution process were identified and their contributions to different libraries compared. In addition to this general characterization of libraries and the deconvolution process, ASMv1.0 offers a means to filter the deconvolution data of screen result for those variants expected by design, or to identify high abundance variants independent of the library design. The results of these analyses have informed the choice of strategies for saturation mutagenesis at multiple steps including library design, screen setup, and data analysis. We believe that the analytical principle and screen strategy presented here, along with our library designer, variant-calling software, and data analysis tools will help enable the scientific community to fully utilize this technology in research applications.

Although, as noted, producing the desired amino acid changes using 2 or 3-nt changes versus the reference sequence is superior to using 1-nt changes versus the reference sequence, it should be noted that the strong hits among the 1-nt delta variants may still score well, even with some suppression of the apparent log-fold change results due to miscalls of reference sequence that match the 1-nt variant. The effect of miscalls on 1-nt delta variant counts becomes less of an issue for short ORFs (e.g., *KRAS*). As illustrated in **Supplementary Figure 6**, the longer the ORF, the more copies of the reference codon that will undergo PCR-based amplification and NGS, and the more copies of each actual designed 1-nt variant that will be also read out, and consequently also the bigger the ratio of the 1-nt delta miscalls to correct calls for the 1-nt variant molecules truly present. To better interpret any 1-nt delta codons that in a variant library, we recommend adding the clonal template ORF as a control sample and processing it in parallel with the screening samples. This control sample may be used as a count-correction factor (as shown in **Fig. 4b**, right) or simply to flag those 1-nt delta variant sequence that occur at high rates due to miscalls of the reference sequence, contaminating and therefore suppressing any fold-changes of that variant between screen arms.

The recent establishment of an open-source platform, MaveDB, to enable variant function data sharing, is a significant development [23] and is a good step towards more widespread sharing of saturation mutagenesis data, reagents and resources. A well-designed variant library should be useful for many different screens of that same genes, provided that the choice of reference sequence upon which the design is based is satisfactory for each application. For example, Boettcher *et al* [24], used the existing *TP53* site-saturated variant library created by Giacomelli *et al* [9], to demonstrate that the dominant-negative activity of *TP53* missense mutations in DNA binding domain was selected-for in myeloid malignancies. A shared inventory of pooled gene variant libraries would be a powerful tool for variant-to-function mapping and common practices for the design composition and readout of these libraries will enhance their utility.

### Conclusions

Complex ORF variant libraries with all possible single-amino-acid substitutions are powerful tools that enable one to exhaustively test all possible protein variants simultaneously in a single experiment. Each experiment produces extensive dictionary of variants and functions. This technology has high access barriers as it demands expertise across many disciplines, and the costs of producing and utilizing the libraries are large. This report explores some factors that are important for screening very large variant pools efficiently and presents a set of well-tested practices and tools to conduct pooled saturation mutagenesis screens.

## Methods

Experimental methods for the published data that were employed in this study are available in the original publications. Nonetheless, for convenience we provide again here methods for all the experimental steps of large-pool variant screening that incorporate our current best practices, based on experience from over 20 variant screens (**Supplementary Table 1**) along with the analysis and computational methods and the data-driven evaluations of library compositions and variant screen error modes that are presented in this study.

### ORF expression vector

A lentiviral delivery expression vector that we have frequently used to clone site-saturated variant libraries is pMT025 (**Supplementary Fig. 7**). This vector was used for the libraries and screens that are re-analyzed in this report and for numerous other ORF variant libraries (**Supplementary Table 1)**, It is suited for restriction/ligation cloning using NheI/BamHI, or NheI/MluI restriction enzyme pairs. The vector has a puromycin resistance selection marker driven by an SV40 promoter. After the cloning, the ORF expression is driven by EF1α. To facilitate cloning, the ORF sequences are flanked with constant sequences carrying the highlighted restriction enzyme sites for cloning:

GGTTCAAAGTTTTTTTCTTCCATTTCAGGTGTCGTGAG**<u>GCTAGC</u>**GCCACC [ORF with ATG start and stop]**<u>GGATCC</u>**CGGGACTAGT**<u>ACGCGT</u>**TAAGTCGACAATC

Two different library construction techniques are described here, one in which the ORF variant sequences originate as pooled double-stranded DNA fragments encompassing the entire ORF, purchased from Twist Bioscience. For the other method, the variants are assembled per ‘tile’ from a pool of single-stranded DNA oligonucleotides to create a plasmid DNA pool of the variants in a pUC57 vector. For the two methods, the resulting variant expression libraries, the screening, and data processing of the libraries are the same, but these two methods demand some differences in library design and library cloning.

### Choice of reference sequence and the template ORF plasmid

An ORF variant library is built from a common template “reference” sequence to which mutations are introduced. This reference sequence may be some version of the wild-type coding sequence of a gene, or it may itself already possess known functional mutations to which additional mutations are added to form the library [25]. In this paper, we call this template sequence the “reference sequence.” It is important at the outset of library design to ensure that the chosen reference sequence will be fully compatible with all the requirements of library construction and screening. For example, in the cases where a preliminary choice of ORF reference sequence contains restriction sites needed for the library cloning, the reference sequence needs to be modified with silent mutations to remove these sites. If the screening assay has been developed using expression of the initial reference sequence, then any changes to the reference sequence should be supported by experiments to demonstrate that these changes have no impact on the reference sequence phenotype and assay. While silent mutations to the should typically induce no change in phenotype, they do have the potential to possibly alter expression (e.g. transcript stability, translational efficiency, etc) and therefore phenotype. In general, we recommend that, prior to library construction, the screening assay be fully vetted using the *exact* reference sequence and expression vector that the library will employ, even if that requires repeating assay development experiments that were previously performed using a similar vector or ORF sequence. It is equally important that a clonal plasmid of the template ORF is cloned into the vector, made into virus, and tested for transduction, selection conditions and genomic DNA preparation efficiency and quality. Finally, the genomic DNA should be, tested to determine the PCR conditions that amplify the ORF with high efficiency since this will need to be done at scale later for the screen deconvolution. Twist Bioscience uses the clonal reference sequence plasmid as the PCR template to produce the mutagenesis library.

### Library design

A saturation mutagenesis designer written in R is available (see the links listed in **Supplementary Table 2**). The reference ORF sequence is first checked to ensure that the restriction sites to be used in cloning are excluded, and otherwise the sequence must be modified with silent changes to eliminate any occurrences of the cloning sites. NheI/BamHI and NheI/MluI cloning sites have been successfully employed for numerous ORF libraries. The designer algorithm steps from N-terminus to C-terminus of the template ORF one amino acid position at a time to select variant codons. For a given template codon there are 63 possible variant codons from which to choose for each amino acid position. These 63 codons are divided into two bins: 54 codons that have 2- or 3-nucleotide differences relative to the template codon (2-3-nt delta), and 9 codons with a single-nucleotide difference (1-nt delta). The codons that would violate the restriction sites exclusion rule are removed from both bins. To design each missense or nonsense variant for each amino acid position, we first select from the 2-3-nt delta codon bin, and then in cases for which no 2 or 3-nt delta option is available, from the 1-nt delta codon bin. If within a bin there are multiple codon choices available for an amino acid variant, the codon is picked by sampling the available codons with probability weightings according to the codon usage frequency of the organism (*Homo sapiens* in all our experiments to date, http://www.kazusa.or.jp/codon/cgi-bin/showcodon.cgi?species=9606). For silent codon changes we pick only from 2-3-nt delta bin and do not include any silent change variant in the library for those residues for which no 2 or 3-nt delta option is available. In the end, each of the designed variant codons substitutes for its corresponding template codon in the full-length reference ORF as one member of the library.

### Mutagenized ORF Library cloning

Saturation mutagenesis library production requires that variant-defining codons be encoded in chemically synthesized DNA oligo strands. For the MITE method, once the 150-nt DNA oligo strands bearing the mutagenized codons are produced, they are first amplified by PCR and then cloned into a high copy number entry vector, pUC57. For Twist Bioscience method, Twist uses the mutant DNA oligo strands as PCR primers to introduce codon changes into the full-length ORF template to produce a pool of properly flanked and variant-carrying full-length ORFs that are ready for cloning into pMT025 ORF expression vector.

#### MITE method

The region of the open reading frame to be mutated is partitioned into 90-base tiles. Oligos representing all mutations in the space of a tile were synthesized. Each 90 base tile is flanked by 30 base sequences that are complementary to the reference ORF. Oligos for adjacent tiles should be synthesized on separate arrays. Oligo tile pools were amplified by emulsion PCR using primers to the 30 base flanking regions and purified on a 2% agarose gel. Entry vector cloning was performed on a per tile basis by linearizing the entry vector at the appropriate tile position using Phusion polymerase (New England Biolabs) and primers to the regions flanking the tiles. The linearized vectors were purified on 1% agarose gel and DpnI treated. The DpnI-treated linear plasmid backbones were then mixed with the relevant PCR amplified tile and assembled by *in vitro* recombination with the NEBuilder HiFi DNA Assembly Master Mix (New England BioLabs). The assembly reactions were purified, electroporated into TG1 *E. coli* cells (Lucigen), and recovered for one hour at 37°C in Recovery Media (Lucigen). Aliquots from the transformations were used to inoculate overnight cultures of LB containing 25 μg/mL of Zeocin (ThermoFisher). Cells were harvested by centrifugation and plasmid DNA was isolated using the QIAGEN Midiprep Plus kit (QIAGEN). Plasmid pools, each corresponding to a tile, were verified by sequencing Nextera XT (Illumina). A pool of all pUC57-tile libraries was made with equal weight per variant. This pool was subjected to restriction digest (e.g., NheI and BamHI). After purification by 1% agarose gel, the excised linear fragment library is ready for cloning ligation with an expression vector that was pre-processed with the compatible restriction enzyme pair.

#### Twist Bioscience method

The full-length variant pool made by Twist Bioscience was delivered as linear fragments of the full-length ORF flanked with adapters for cloning. This fragment library was first digested with NheI and BamHI overnight, and then underwent ligation with pMT025 vector pre-processed with the same restriction enzyme pair.

#### Cloning ligation

For either cloning method, the ligation was carried out with 5:1 insert-to-vector molar ratio, using T7 DNA ligase at room temperature for 2 hours. The ligation was cleaned up with isopropanol precipitation. The resulting DNA pellet was used to transform Stbl4 bacterial cells. Plasmid DNA (pDNA) was extracted from the harvested colonies using QIAGEN Maxi Prep Kits. The resulting pDNA library was sequenced via the Illumina Nextera XT platform to determine the distribution of variants within the library.

### Virus preparation and titer measurement

Lentivirus was produced and quantified using the protocol available at http://www.broadinstitute.org/rnai/public/resources/protocols. Briefly, 293T viral packaging cells were transfected using TransIT-LT1 transfection reagent (Mirus Bio) with the pDNA library, a packaging plasmid containing *gag, pol* and *rev* genes (psPAX2, Addgene), and an envelope plasmid containing VSV-G (pMD2.G, Addgene). Media was changed 6-8 hours after transfection, and virus was harvested 24 hours thereafter. The virus titer was measured by infecting 3000 A549 cells per well in 96-w cell culture plate with a series of diluted standard virus of known titer for a standard curve, and a series of diluted library virus of unknown titer to find data points that lie in the linear range of the standard curve. Twenty-four hours post infection, the cells were subjected to puromycin selection for 2 days. Cell number/survival post-selection was quantified with alamarBlue.

### Determine cell numbers required for library representation

The number of cells transduced per replicate and the number of cells seeded per replicate at each passage was typically chosen to be an average of 1000 cells per variant in the library pool. This minimum cell number was then preserved throughout cell culture passaging. For positive selection screens, the majority of the cell population is depleted upon introduction of the selection pressure and only the subset of cells that carry survival-conferring variants will survive. In these cases, adequate cell number representation of the full library must be maintained until the selection pressure is applied, but as the selection pressure takes effect, the cell numbers are reduced, and the resulting smaller cell population, enriched for survival-conferring variants is collected at from these selected arms at the end of the screen. For negative selection screens, the majority of the library will be present at the end of the screen, and the end-point samples should comprise the same number of cells, e.g. 1000 cells per variant (i.e., per library member), that was maintained throughout.

### Genomic DNA preparation

Upon the completion of the screen, cell pellets were collected. Genomic DNA was extracted from cell pellets using the Machery Nagel NucleoSpin Blood Kit (Midi, Clontech Cat. #740954.20; XL Maxi, Clontech Cat. #740950). The detailed protocol is available at http://www.broadinstitute.org/rnai/public/resources/protocols.

### Determine the number of PCR reactions per sample

For positive screen end-point samples, the number of cells that need to go through ORF-extracting PCR is determined in such a way that for each library variant, gDNA template from 500 or more cells will be amplified by PCR. For reference samples and negative selection screen end-point samples, the cells/variant ratio should be maintained at 1000 or more. PCR is sensitive to the amount of gDNA input and prior to screening, these conditions must be optimized. For example, in 50ul PCR reactions, we test different amounts of gDNA, using, for example, a series of 250 ng, 500 ng, 1 μg, 2 μg. This titration will determine the optimal concentration of gDNA that can be used to produce robust PCR products. The gDNA loaded into each PCR reaction should not exceed the amount that can be successfully amplified for the reaction conditions and scale employed. Diploid human cell lines have roughly 6.6 pg DNA per cell. Here is the formula to determine how many PCR reactions are needed for a sample, again making sure not to exceed the maximum ng of gDNA per well that can be amplified:

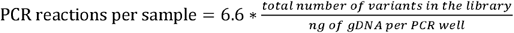

### Design PCR primers for ORF amplification from gDNA by PCR

The ORF sequence is amplified from the genomic DNA by PCR. The PCR products are randomly sheared with a transposon insertion method, index labeled, and sequenced with next-generation sequencing technology. Each sequenced fragment is generated by 2 transposon insertion events so that both ends of the fragment are tagged with a transposon. Therefore, the ends of the initial gDNA PCR product will be less frequently represented in the NGS library and will have shallow coverage. For this reason, we advise that the PCR primers be designed include an extra ~100 bp of sequence at each end beyond the region of ORF that is mutated in the library. For the pMT025 vector carrying libraries whose variants span entire ORF insert, we use these 2 primers:

Forward: 5 ‘-ATTCTCCTTGGAATTTGCCCTT-3’
Reverse: 5 ‘-CATAGCGTAAAAGGAGCAACA-3 ‘

### ORF amplification from gDNA by PCR

PCR reactions were set up in 96-well plates according to the optimized PCR condition and number of PCR reactions per sample calculated above. The volume of each PCR reaction was 50 μL and contained an optimized amount of gDNA. Q5 DNA polymerase (New England Biolabs) was used as the DNA polymerase. All PCR reactions for each gDNA sample were pooled, concentrated with a PCR cleanup kit (QIAGEN), and separated by gel electrophoresis. Bands of the expected size were excised, and DNA was purified, first using a QIAquick kit (QIAGEN), and then an AMPure XP kit (Beckman Coulter).

### Nextera sequencing

Nextera reactions were performed according to the Illumina Nextera XT protocol. Each sample was split into 4 Nextera reactions, each with 1 ng of purified ORF PCR products. Each reaction was indexed with unique i7/i5 index pairs. After the limited-cycle PCR step, the Nextera reactions were purified with AMPure XP kit. All samples were then pooled and sequenced using an Illumina sequencing platform chosen among Miseq, Hiseq, Nextseq, or NovaSeq, according to the number of reads needed as calculated below.

### Determine number of reads needed for the screen

The Nextera-produced fragment molecules must be sequenced at adequate depth to represent every variant. The number of total reads required depends on ORF length, total number of variants in the library, read length, read quality etc. The minimum number of read required can be estimated as follows:

For negative selection screen samples, or reference samples:

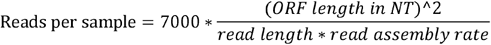

Where *‘read length’* is typically 300, as we use paired-end sequencing reads, 150 bases per read, and *‘read assembly rate ‘* is in range of 0.7-0.9.

For a positive selection screen sample:

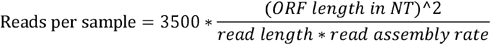

### Determine the optimal parameters for variant-calling software ASMv1.0

NGS is performed on transposon-sheared fragments which are not uniform in size. The Nextera conditions as recommended typically achieve an average fragment size of approximately 140 base pairs, data not shown). For fragments longer than 2x read length (e.g., 2 x 150 = 300 bases), the pair of reads are disjoint (non-overlapping); for fragments shorter than the read length (e.g., 150 bases), the reads are fully overlapping, and each extends into the transposon adaptor located in the opposite end; for fragments sized between 1x and 2x read length (e.g., 150-300 bases), the pair of reads overlap but do not extend into the transposon sequences. The variant calling software first aligns reads to the reference sequence. The transposon tails are then clipped off at the coordinate determined by the start of its paired read. It then applies the ***min-q*** (minimum base quality) filter to trim the end of the aligned reads to ensure there are at least 15 bases from the trimmed end inwards with base quality above the min-q parameter. The ***min-read-length*** filter admits the trimmed reads that meet the selected length threshold. All failed reads go into the low-q bin. Next, the filter of ***min-flank-length*** is applied that requires reference bases at end of the trimmed reads. A min-flank-length of 2 or more bases excludes cases in which an altered codon is split between 2 Nextera fragments. It also prevents erroneous variant calls due to reads that end with transposon sequences for which transposon clipping failed to work due to a lack of paired read overlap. Finally, a variant is called only when the underlying base changes are above the min-q threshold.

To tune our software parameters, we used the sequencing data of a saturation mutagenesis plasmid DNA library for *PDE3A* gene. This library was built by mutagenizing a region of 1425 base pairs, using a variant oligonucleotide pool produced by Twist Bioscience. The errors in the library are concentrated in within 30 bases of a planned codon change (**Fig. 3**). These errors can be explained by mutagenesis primer synthesis errors. The codon changes are located in the middle of the mutagenesis primers, sandwiched between two 20-30 base pair flanks. In next-generation sequencing, the transposon sheared fragments are, on average, 140 base pairs, similar to the sequencing read length. Characteristic to the way how the variant libraries were synthesized, the close proximity of codon modifications, including the planned and the unintended, may indicate that these codon modifications reside on the same transposon-sheared fragment and on the same sequencing read. Therefore, each library ORF, after transposon shearing, produces one variant-bearing fragment; the rest of the fragments are all wild-type. That is, if we ignore the miscalls introduced by PCR, the wild-type/variant read ratio of this library should be: 1425/140 −1 = 9.2. This ratio, 9.2, represents the highest ratio we could observe, as any PCR miscalls would only shift this ratio lower.

In determining the PCR miscall rate, we processed a clonal plasmid of pMT025-PDE3A. The sequencing of this clonal plasmid DNA, which had undergone the same PCR process as the saturation mutagenesis plasmid DNA library, showed error rate of 0.00037 miscalls/sequenced nucleotide.

Start with miscall-free wild-type/variant read count ratio 9.2:1, At this miscalls rate, one expects fewer reference reads versus variant reads, namely (9.2*140*0.00037)=0.48 of the expected reference reads incur miscalls and appear as variants, predicting a wild-type/variant read count ratio:

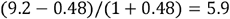

By processing the same plasmid DNA library sequencing data set with various parameters, we were able to identify the optimal parameters that produce a wild-type/variant ratio of 5.9, and yield reasonable output of both wild-type and variants: min-q = 30, min-read-length = 40, min-flank-length = 2.

### NGS Data processing and analysis

The data from the NGS pipeline are in .fastq format. Each NGS cluster produced 2 long reads (150 bases each), and 2 index reads (each 8 bases long). The 2 indexes were used to label the screen samples, so that many samples can be pooled and sequenced together and the reads that originating from each sample can then be distinguished in the sequencing data by these indices. Site-saturation variant library screen data file sizes are typically 4-5TB. Generous computer storage is needed to process the data, especially as the reads go through the filtering process (**Supplementary Fig. 2**) prior to variant calling.

#### Read filtering

The reads from an NGS fragment are trimmed (1) by base quality, (2) by removing non-ORF transposon sequences, (3) by requiring that mutated nucleotides are flanked by, at minimum, the number of reference-matching bases that is specified by the min-flank-length parameter, and (4) by requiring the trimmed reads be no shorter than the min-read-length (see **Supplementary Fig. 2**).

#### Variant calling

There are two versions of variant calling software, ORFCallv1.0, the earlier version; and ASMv1.0, the current version. The reads that have passed the filtering step are processed differently by ORFCallv1.0 and ASMv1.0:

i. ORFCallv1.0 (an earlier version software; Giacomelli *et al* [9]): Variants are tallied in the context of a single codon space. Without regard of the association of the multiple changes physically in a single molecule, ORFCallv1.0 essentially scans from N- to C-termini, one amino acid position at a time, and sums up the occurrence of codons or amino acids at each position.
ii. ASMv1.0: This is a molecule-centric approach that is more sophisticated than the approach of ORFCallv1.0. The NGS dataset is analyzed in its entirety, with the following new capabilities: (1) The trimmed reads are evaluated full-length; that is, ariants are called in the context of entire read (or read pair) and by nucleotides, not amino acids. The programmed variant is called when 2 conditions are met: (i) a library-designed nucleotide change are detected (in 1-3 nucleotides within a codon position), and (ii) there are zero ***additional*** nucleotide variations throughout the entire read or read pair. (2) Indel detection: This feature was lacking in ORFCallv1.0. This new feature of ASMv1.0 allows us to process libraries that include indel variants. In ASMv1.0, mutations in a molecule are depicted at the nucleotide level by position along the entire pre-framentation PCR product. They are annotated by listing positions of change followed by the change itself. For example, ‘a change from G to A at position 1140’ is described as ‘**1140:G>A’;**‘a deletion of an A at position 1141’ is ‘**1141:A>-**’; ‘an insertion of a T after position 1142’ is ‘**1143:->T**’. Also, with ASMv1.0, a host of annotations formatted as a datafile can be derived from the variant descriptions. For example, a *TP53* variant with description ‘**1140:G>A, 1141:A>C, 1142:G>C**’ is annotated as (a pair of brackets marks an annotation): [E343T] [Intended] [one codon change] [343:GAG>ACC] [missense] [substitution]. Another *TP53* variant with description ‘**1140:G>A, 1141:A>C, 1142:G>C, 1145:G>T**’ is: [E343T] [Unintended] [two codon changes] [343:GAG>ACC, 344:CTG>CTT] [substitution] [missense/nonsense] etc. These two variants encode the same amino-acid variant. The first variant is an intended variant that matches a designed nucleotide-variant of the library. The second variant is the same amino acid call as the first variant, but it is annotated, rightfully, as an unintended variant carrying an extra nucleotide change 4-nucleotide away from the intended codon change - this extra nucleotide change results in a silent codon change.

#### Parsing data into a form ready for hit calling

The output files from the above variant-calling software are still typically quite large in size (over 100MB), large in numbers of files, and not readily useable. These output files need to be parsed, aggregated and annotated. For this purpose, we built a parser called ASMv1.0_Parser, which was written in R, and is available for download (see the links listed in **Supplementary Table 2**). The Parser Pak consists of a ‘readme’ file detailing the instructions, the scripts (in R), and some training data sets. Through the parser, all data files are organized into a single .csv file, one column per screen sample. These .csv files include a short version that is a subset of the planned variants only, and a complete version that includes all called variants, including planned/unplanned, insertion/deletion/substitution, etc. In addition, we also report the ORFCallv1.0 version of data.

#### Scaling

We use read counts from the NGS data as the quantitative measurements of variants. Reads that entirely match the reference sequence are not informative, since they are presumably all reflect the unmutated, uninteresting parts of an amplified ORF molecule (the majority) that did NOT contain the mutation that defined that specific ORF variant. Yet these uninformative reads are important for scaling the variant counts, as a way to mitigate the transposon insertion biases. A called variant, either intended or unexpected, is defined by all nucleotide variations (these variations together are the variant signature that define the variant) detected between a read pair and the reference sequence. The tallied counts of a given variant signature may fluctuate depending on the transposon insertion bias in ORF DNA fragmentation step as depicted in **Supplementary Figure 9**. However, we can remove these biases by scaling the variant count against the total coverage of reads, both wild-type reads and variation-bearing reads, which span the full region that is marked by the variant signature; this read coverage is reported in ASMv1.0 output as a separate column called ‘reference_count’ next to the ‘count’ column. While all molecules in a library are full-length ORF and at each nucleotide position, one may expect uniform depth, we have demonstrated that this is not the case (**Supplementary Fig. 9**). To determine the relative abundance among variants in the same sample, as in **Supplementary Figure 10**, the raw counts should first be scaled to the fraction of the total read coverage across the variant signature. However, in cases that compare the variant abundance change **between** two samples, such as computing fold-changes in screening samples, scaling the raw counts may not be necessary, as the positional raw count variations caused by transposon insertion bias will cancel out between two samples when computing fold-changes (**Supplementary Fig. 11**).

## Acknowledgements

We thank our collaborators of saturation mutagenesis projects, Alice Berger, Jesse Boehm, Ben Ebert, Laura Evans, Eejung Kim, Shengwu Liu, Julie-Aurore Losman, Huong Ly, Peter Miller, Utthara Nayar, and Nikhil Wagle for demonstrating the diverse utilities of the technology, Broad Institute Genomics Platform for providing NGS support, Nancy Tran and Cong Zhu for the early work on saturation mutagenesis method, Stephanie Milczarek, Kirsten Milenkowic, Michael Zagieboylo, and Jonah Schwartz for construction of some of the variant libraries referenced in this study, Leon Yang for critical reading of the manuscript, and to all the members of Broad’s Genetic Perturbation Platform for their help and support. This work was supported by NCI U01 CA176058 (WCH), the Mark Foundation (WCH), American Cancer Society Professorship (MM), American Cancer Society Mentored Research Scholar Grant MRSG-18-202-01 (ALH), Department of Defense W81XWH-18-PRCRP-CDA (ALH), Team Lick Cancer (ALH), Unravel Foundation (ALH), and Wong Family Award (ALH), and the Functional Genomics Consortium (D.E.R.).

## Author Contributions

X.Y., D.E.R., A.L.H., A.O.G., and T.S. wrote the manuscript. W.C.H., M.M., H.G., A.J.A., R.E.L., N.P, M.M, A.B., and F.P. reviewed and edited the manuscript. A.L.H., X.Y., T.S, and D.E.R. designed the new variant-calling software ASMv1.0. T.S. wrote the variant-calling software ORFCallv1.0 and ASMv1.0. X.Y., A.L.H., D.E.R., and T.H. optimized the run parameters of ASMv1.0. X.Y. wrote R codes to design site-saturated variant libraries and to parse the output of the variant-calling software. X.Y. conducted data analysis with input from D.E.R., A.L.H., A.O.G., and assistance from B.P.L. D.A and T.G. adapted the variant-calling software into a saturation mutagenesis data pipeline. X.Y., A.L.H., A.O.G, R.E.L., B.P.L., B.F, H.K., E.S., F.P., and L.S. designed and conducted wet-lab experiments. All authors contributed to and approved the manuscript.

