## Supplementary_Tables1-3 for "Defining protein variant functions using high-complexity mutagenesis libraries and enhanced mutant detection software ASMv1.0"

**Supplementary Table 1: ORF variant libraries and screens**

| Genes | Number of screens processed |
| --- | --- |
| Full_length ORF variant libraries: |  |
| *BFL1* | 1 |
| *DHODH* | 1 |
| *EGFR* | 2 |
| *EZH2* | 0 |
| *FDX1* | 1 |
| *FGFR2* | 0 |
| *HER2* | 0 |
| *IDH1* | 1 |
| *IDH2* | 1 |
| *KRAS* | 1 |
| *KRAS G12C* | 1 |
| *KRAS G12D* | 2 |
| *MCL1* | 1 |
| *SHOC2* | 1 |
| *SMARCB1* | 1 |
| *TP53* | 2 |
| *RHO* | 0 |
| Partial ORF variant libraries: |  |
| *EGFR indel* | 0 |
| *EGFR mini* | 1 |
| *EGFR*_D770_N771insSVD | 0 |
| *EGFR*_E746_A750del | 0 |
| *EGFR*_E746_A750del_T790M | 0 |
| *EGFR*_L858R | 0 |
| *KRAS mini* | 1 |
| *PDE3A* | 1 |
| *PPM1D* | 1 |
| *RIT1 mini* | 1 |
| *SCN2A* | 0 |
| *WRN* | 0 |

**Supplementary Table 2: Download sites of Saturation Mutagenesis tools**

| Software | Download link | Used for | Contact |
| --- | --- | --- | --- |
| ASMv1.0 | As a part of the Genome Analysis Toolkit (GATK v4.2.0.0):  <https://github.com/broadinstitute/gatk/releases> | Advanced variant-calling | Ted Sharpe  |
| ASM_parser.R | <https://github.com/broadinstitute/SatMut_ASM_Parser/releases/tag/v1.9> | Parse ASMv1.0 output files into a single .csv file | |
| VariantLibrary_Designer_v1.9.R | <https://github.com/broadinstitute/SatMut_VariantLibrary_Designer/releases/tag/v1.9> | Pick variant codons using the design rules stated in this study | |

**Supplementary Table 3: The functionality comparison of ORFCallv1.0 and ASMv1.0**

| Variant-calling software | ORFCallv1.0 | ASMv1.0 |
| --- | --- | --- |
| Substitution mutation | Yes | Yes |
| Indel mutation | No | Yes |
| Space defining a variant | 3 nucleotides (1 codon) | Entire sequencing read, or read pair |
| Variant description/annotations | No | Yes |
| Analysis run-time | Hours | Days |
| Library error assessment | No | Yes |
