## Supplementary_Figures1-11 for "Defining protein variant functions using high-complexity mutagenesis libraries and enhanced mutant detection software ASMv1.0"

**Supplementary Figure 1. A saturated ORF variant library that mutagenizes 400 amino acid positions, with 20 codon changes per position.** When all 8000 variant clones are aligned, at each codon position, there are 20 variant-defining codons (in red), and 7980 repeats of the same template codon (in black).

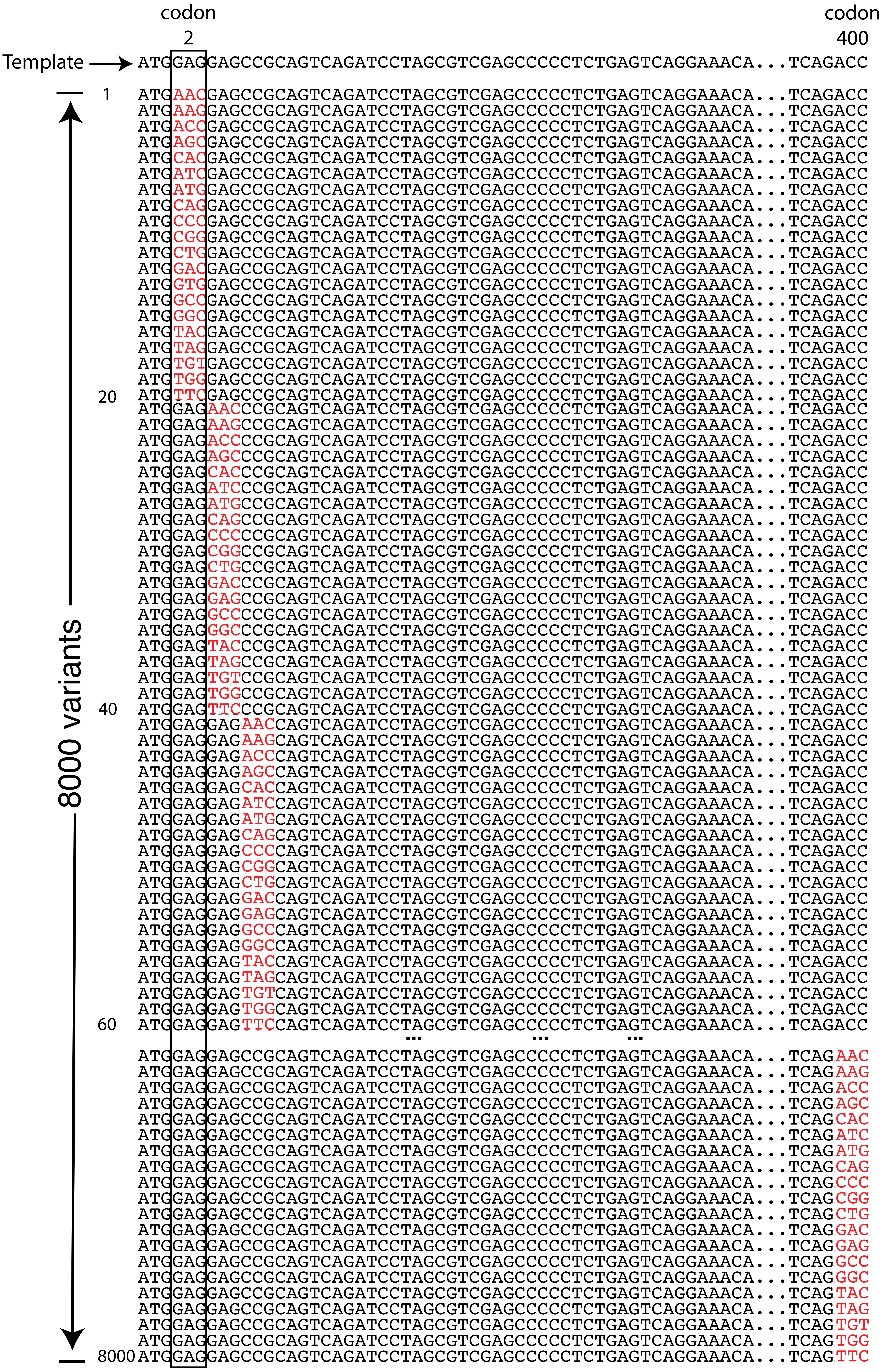

**Supplementary Figure 2 The inner works of the variant calling software ASMv1.0 and the placement of reads.**

#### How the reads are processed

- Align to reference
- Clipping off the transposon tail, using its paired read, in cases that the sheared ORF fragments are shorter than the read length; the start of the paired read marks the end of the sheared ORF fragment
- Trimming ends by min-q filter (optimal: 30)
- Apply min-read-length filter (optimal: 40 bases)
- Apply min-flank-length filter (optimal: 2 bases)
- Call a variant only when the changed bases are above min-q

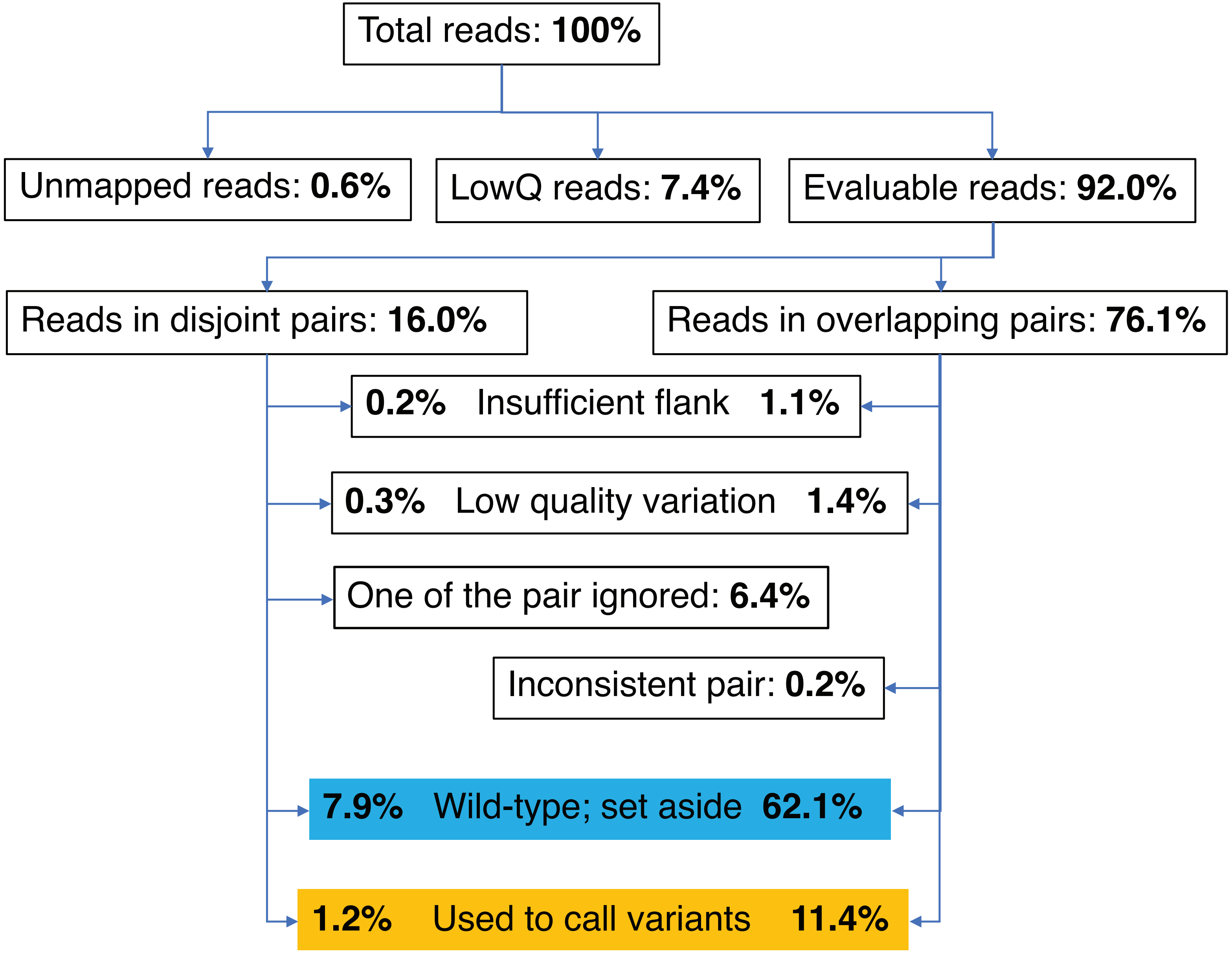

**Supplementary Figure 3. Screen results of p53$^{NULL}$ A549 cells infected with the library, and then treated with etoposide.** Under etoposide selection pressure, the wild type p53 alleles in A549 would promote cell survival. Thus, in this p53$^{NULL}$ A549 cells/etoposide screen, the wild-type-like *TP53* variants are enriched and *TP53* loss of function (LOF) variants are depleted. The arrangement of figure panels in this figure mirrors **Figure 1**.

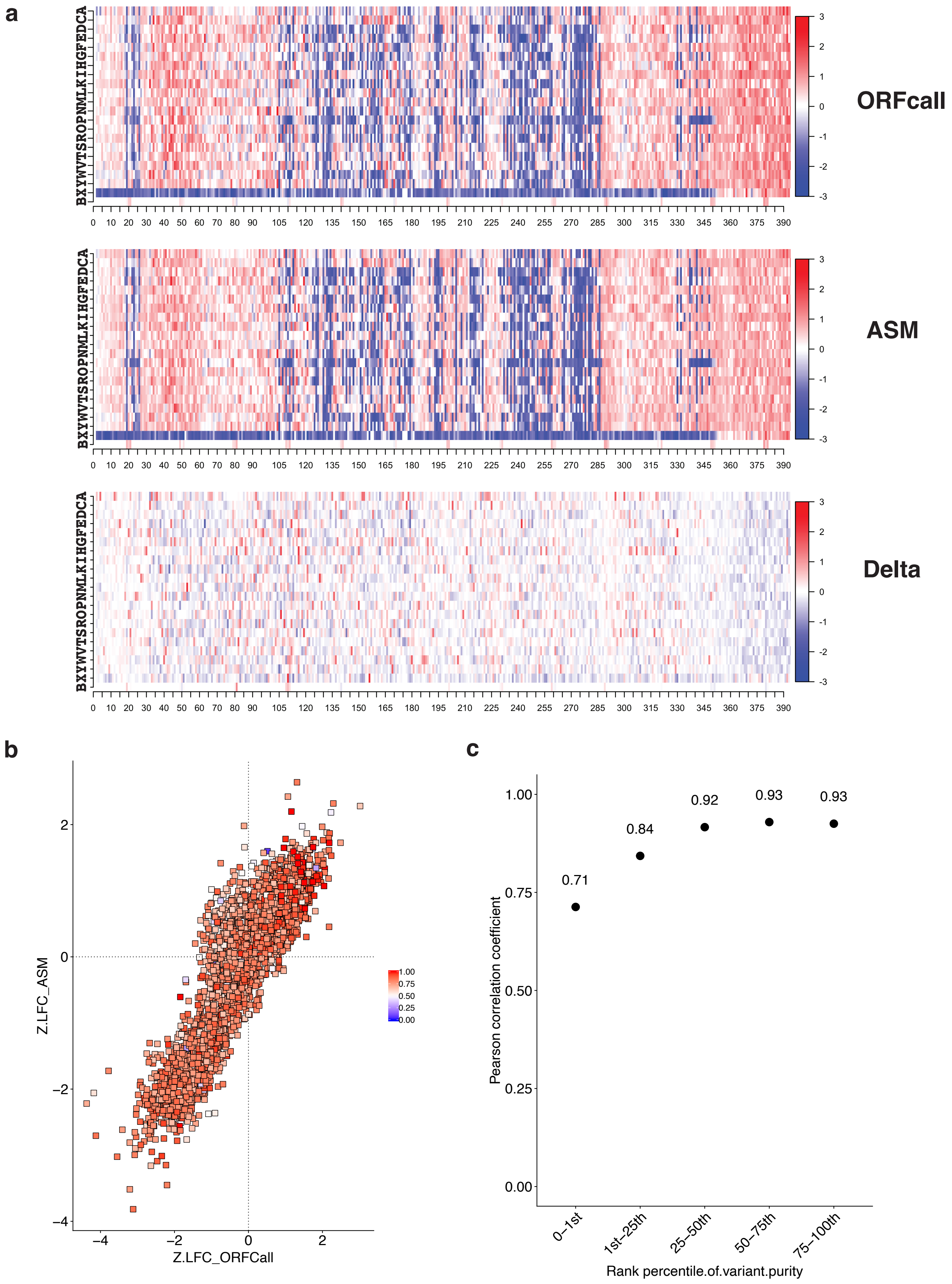

**Supplementary Figure 4. Screen results of p53$^{WT}$ A549 cells infected with the library, and then treated with nutlin-3.** A549 cell line with endogenous wild-type *TP53* (p53$^{WT}$) was infected with the virus particles packaged from the plasmid library. In this screen, under nutlin-3 treatment, which activates p53 by disrupting the interaction between p53 and MDM2, the *TP53* variants exhibiting dominant negative effect (DNE) are enriched. Like **Supplementary Figure 3**, this figure has the same panels arranged the same way as **Figure 1**.

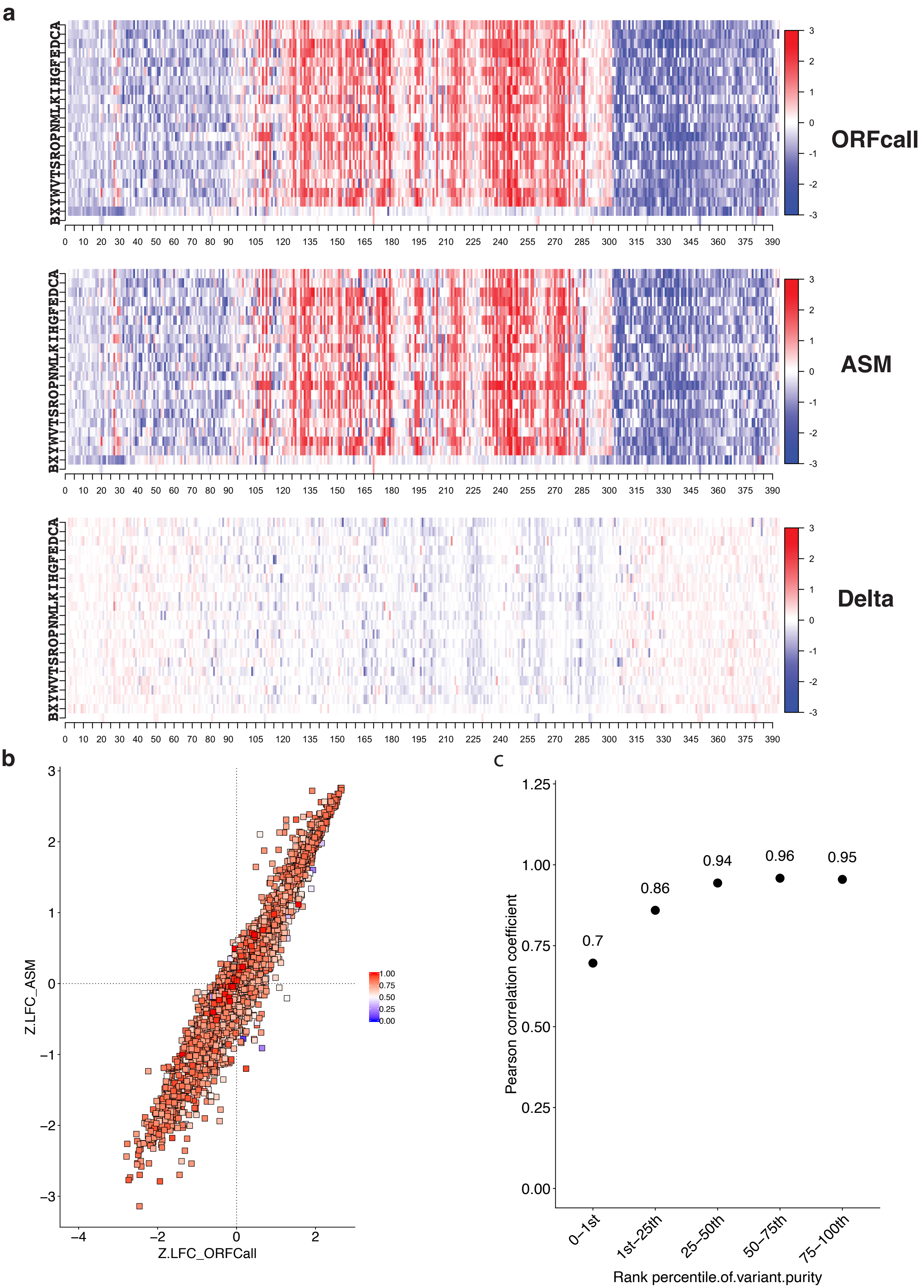

### Supplementary Figure 5

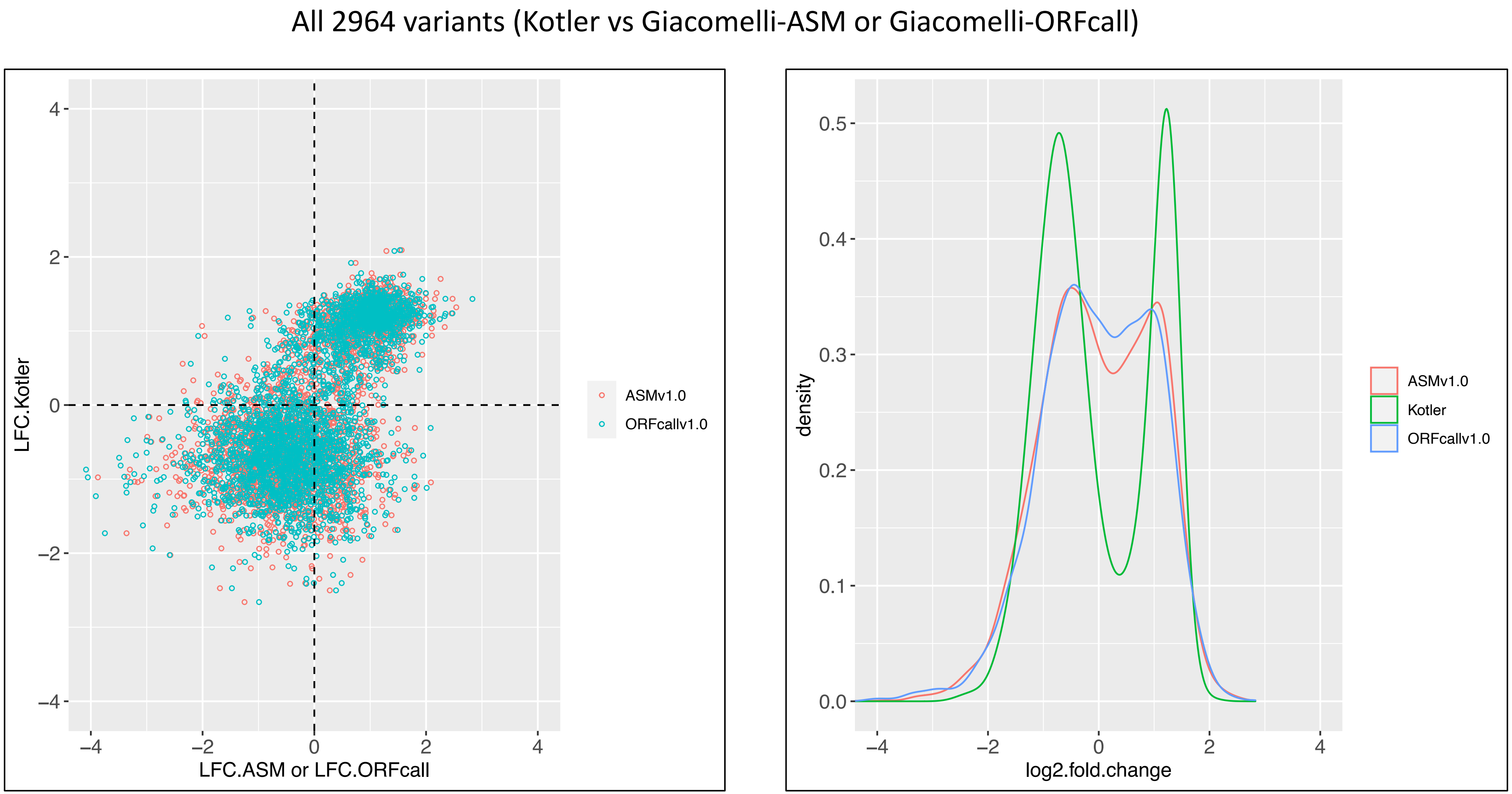


**Supplementary Figure 5. The comparison of the behaviors of shared protein variants in Kotler and Giacomelli screens.** There are 2964 protein variants shared by the two libraries. The scatter plot (**left**) compares the log2-fold-changes in Kotler screen with that of Giacomelli screen called with either software, ASMv1.0 and ORFcallv1.0. Even though the two screens were done with two different libraries, in two different cell lines, and processed with two different variant detection methods, good correlations between Kotler and Giacomelli (called by either ASMv1.0 or ORFcallv1.0) is observed. As pointed out by Kotler *et al* and observed here, all three sets of data demonstrated clean bimodal grouping (**left** and **right**) that marks the p53 loss-of-function (enriched) and p53 wild-type-like (depleted) variants.

**Supplementary Figure 6. Illustration of the effect of PCR errors (miscalls) on variant quantification.**

(a) We consider a hypothetical library for a gene of 400 amino acids in length, with 20 changes per codon position - that is, a library of 8,000 variants. For simplicity, we assume (1) it is a perfect library that has no unintended errors, (2) all variants are evenly distributed, (3) we fully sequence its 8,000 members once each. When all library members are stacked as shown, at each amino acid position, there is a single copy for each of 20 planned variant codons (in red) and 7,980 copies of the reference codon (in black). That is what actually is in the library. In the sequencing readout of the library, however, the PCR error rate in processing the 7,980 copies of the reference codon will result in artifactual codon calls that look like 1-nt delta variants.

(b) At a rate of 0.0003 miscalls per nucleotide readout (the measurements shown in **Fig. 3a**), sequencing 7,980 x 3 = 23,940 reference nucleotides (template sequences) produces 7.2 errors shared among 9 codons (in red), each with a 1-nucleotide deviation from the reference codon. These 7.2 miscalls on average will add 0.8 counts to a planned variant codon that is 1-nt delta in relation to the reference codon. That is, an actual 1-count variant will appear to be a 1.8-count variant. In positive selection screens, a real enrichment from 1 count to 10 counts (10-fold enrichment) looks like from 1+0.8=1.8 counts to 10+0.8=10.8 counts (6-fold enrichment). In drop-out screens, a depletion will never go below the counts of sequencing miscalls, because the depletion cannot go below 0.8 counts from 1+0.8=1.8 counts. That is, a 2.2-fold depletion is the lowest the depletion that can be detected for this 1-nt delta variant.

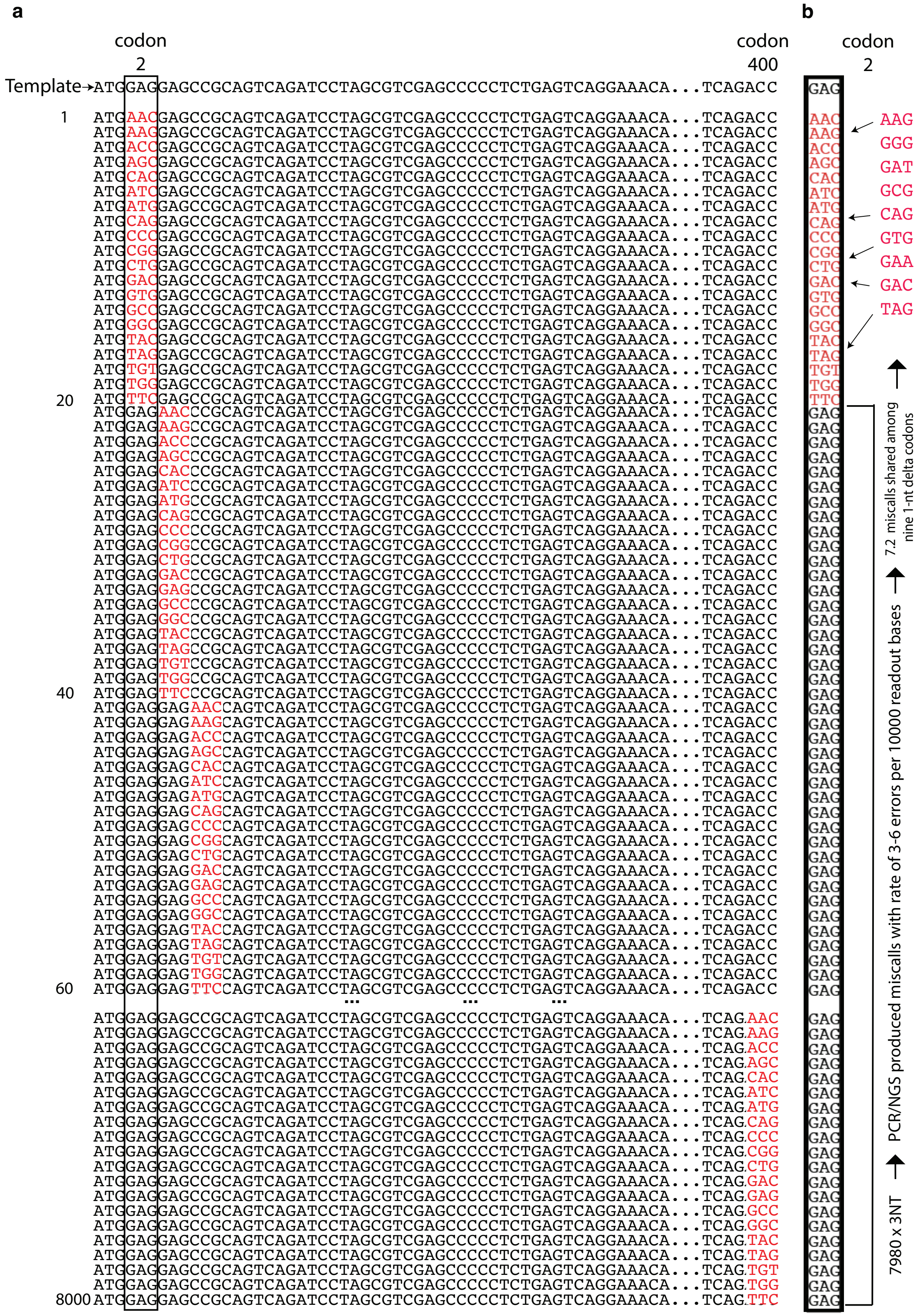

**Supplementary Figure 7. The choices of silent variants.** In most cases, the codons encoding the same amino acids differ at the wobble base of a codon and therefore 1-nt delta silent variants would dominate this control group. In the *SMARCB1* library, we intentionally included nearly all possible silent codons at each position. Similar to 1-nt delta missense variants, the 1-nt silent variants demonstrate inflated counts (a), and artifactually narrowed fold-change range (b), owing to the miscalls introduced by PCR/NGS. We hence reason that the 2-nt and 3-nt delta codons are better choices of silent variants than 1-nt delta codons are.

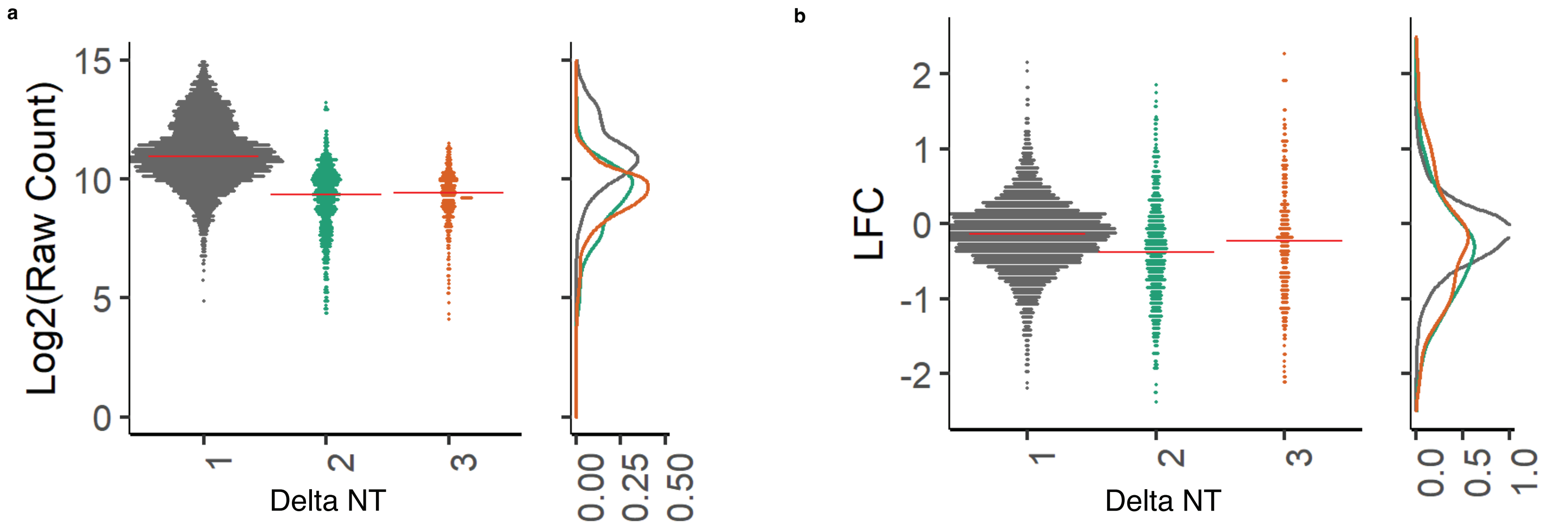

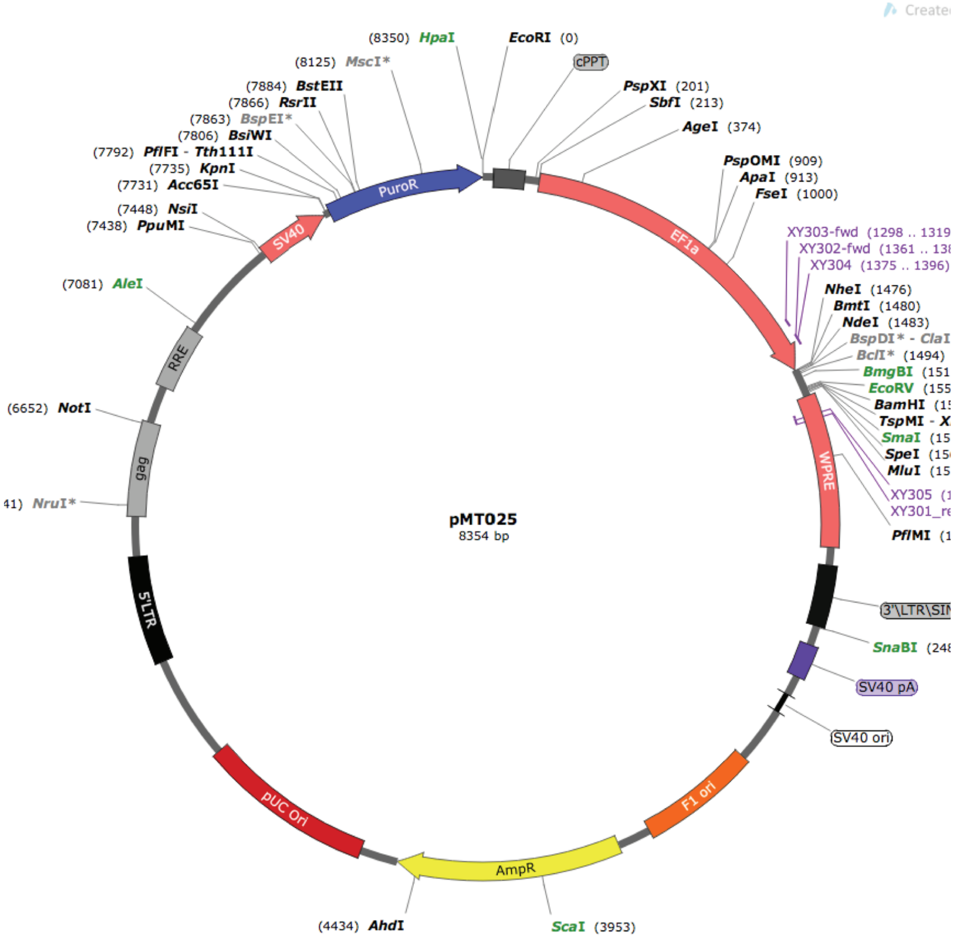


**Supplementary Figure 8. ORF expression vector pMT025.**
This vector was modified from pLX307 vector (https://portals.broadinstitute.org/gpp/public/vector/orfvectorsummary/) by replacing the Gateway cassette with a small stuffer carrying restriction enzyme sites for restriction/ligation cloning. There are two options the pairs of restriction enzymes, NheI/BamHI, or NheI/MluI. In cases that the ORF template has the restriction enzyme sites reserved for cloning, the ORF template will be modified to allow one restriction enzyme pair to be used for cloning. has puromycin resistance selection marker driven by SV40 promoter. After the cloning, the ORF expression is driven by EF1a.

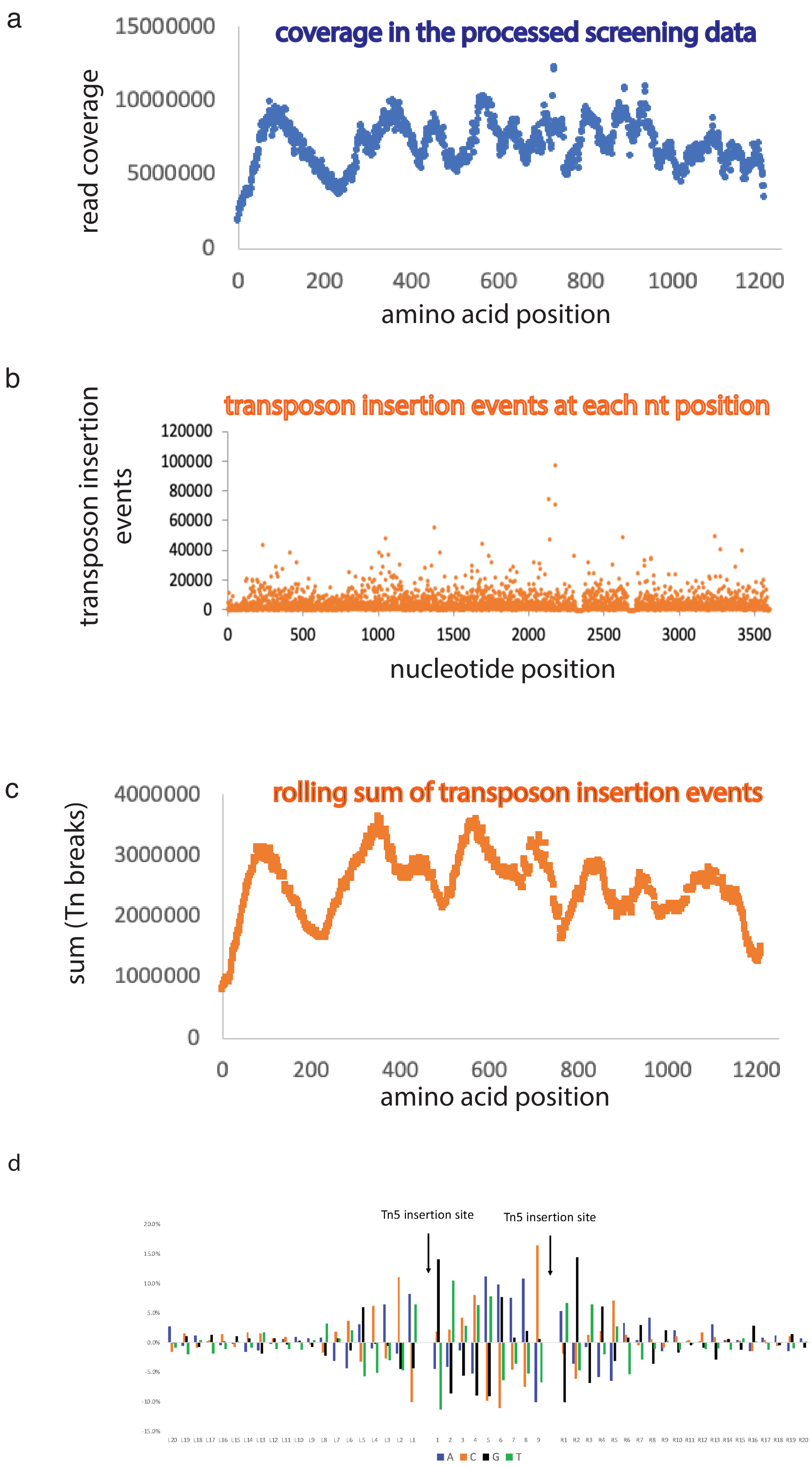


**Supplementary Figure 9. The fluctuation in total coverage along the ORF is explained by transposon insertion bias.** The screen data were generated with the Illumina Nextera pipeline, which includes transposon insertion events to break up the ORF fragments, PCR reactions with indexed primers to bar-code samples, and next-generation sequencing detection. Using the pMT025-EGFR library as an example, the spiky coverage we observed (**a**) is due to transposon insertion bias. We examined the raw sequence for each read pair and located the transposon insertion site. We then summed all insertion events at each nucleotide position (**b**). We tallied the total transposon insertion events of all positions within 140 nucleotides, the average of transposon-sheared fragment sizes, and assigned the value to the center nucleotide position. The resulting sum represents the transposon insertion events that produce reads passing through the center position. After aggregating the sum of transposon breaks as a function of amino acid position, we reproduced a coverage profile (**c**) that mirrors the read coverage profile shown in **a**. Because we conclude that the Nextera shotgun shearing is not random, data scaling, position by position, may become necessary in applications that require comparing the absolute variant abundance, for example, the assessment of library variant distribution. **(d) Tn5 insertion site sequence bias.** In our NGS data, the first bases of a pair of reads mark the transposon insertion sites of a double strand break (DSB), one breaking the top strand, the other bottom strand, and two breaks are 9 bases apart. This allowed us to map out the insertion site preference of Tn5. For all transposition events detected in next-generation sequencing of a clonal plasmid, pMT025-EGFR, the nucleotide compositions (frequencies) at positions relative to the transposon insertion sites are tallied. The nucleotide frequency at each position is subtracted with the nucleotide frequencies of the entire ORF. The adjusted nucleotide frequencies at positions referencing the insertion sites are plotted. We can conclude that the Nextera shot-gun sheering is all but random, as we observe strong sequence biases between (9 bases) and surrounding (5-6 bases each side) the insertion sites.

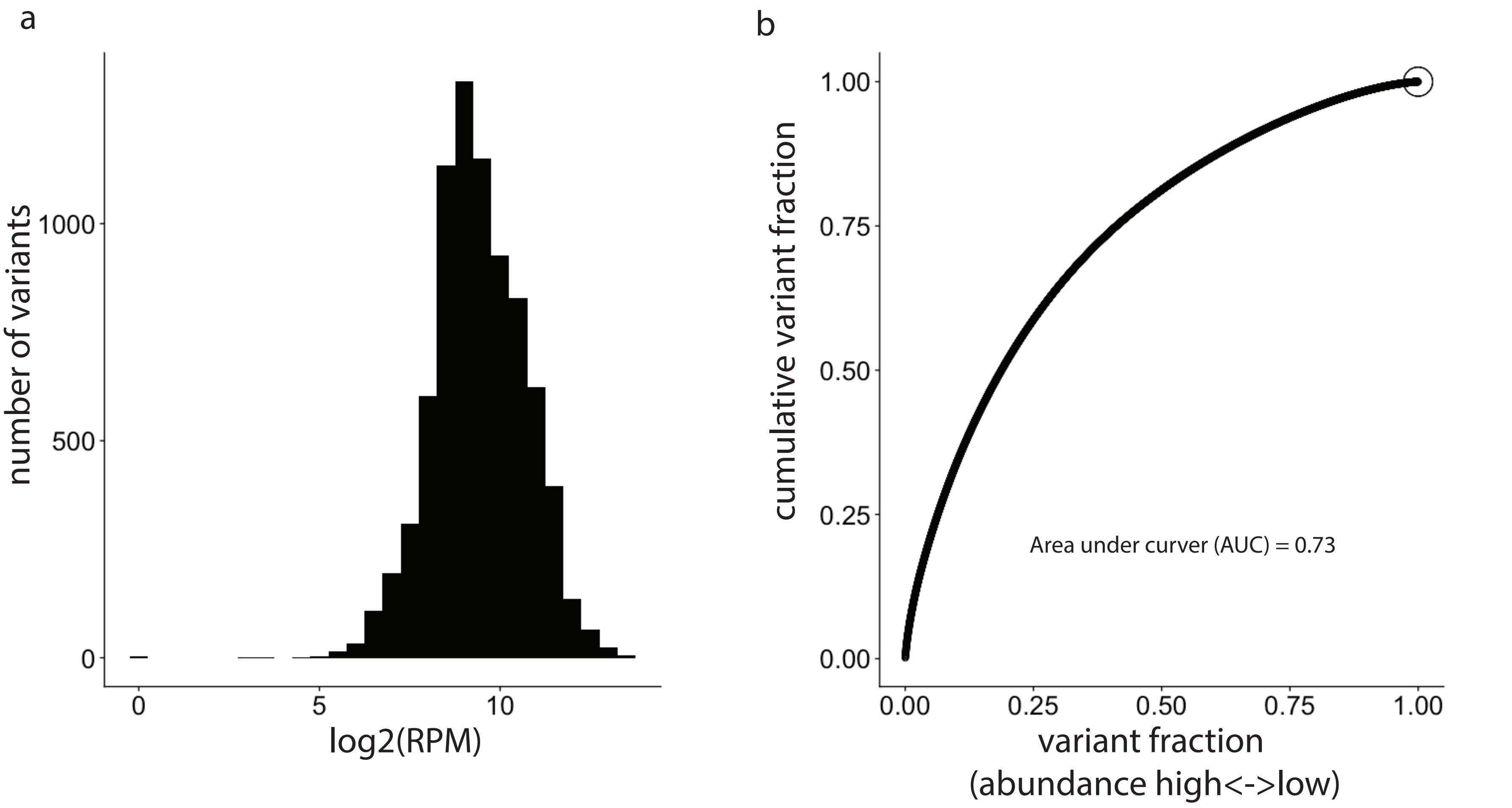


**Supplementary Figure 10. Saturation mutagenesis library variant distribution assessment. (a) Histogram.** We assess clone distribution using scaled abundance. Due to transposon insertion bias, the variant raw counts alone cannot represent the abundance distribution of the members in the library. The raw counts were first scaled to the total read coverage of the same local position, the resulting fractions were then log-transformed into log2(reads per million), which was then used to form the bins of the histogram. **(b) Modified Lorenz curve.** The planned variants were first sorted by the scaled abundance (the fraction of a variant raw count over the position-wise total coverage), in reverse order and placed, one data point for each variant, on x-axis, the cumulative variant scaled-abundance on y-axis; both x- and y-axis are scaled to 1. In the other word, the x-axis is the percentile of library members in reverse (differs from conventional Lorenz curve) order, while y-axis is the cumulative abundance along the x-axis. In this plot, area under curve (AUC) is a good measure of the evenness of library distribution. A 0.5 AUC is for a uniformly distributed library, while an AUC of 1 indicates a case where only 1 variant shows up.

**Supplementary Figure11. Data scaling is not necessary in computing fold-change.** Fold-changes are calculated by comparing 2 samples, for example, a drug-treated sample versus a reference sample. The positional bias in read coverage exist in both samples and the biases may cancel out. With pMT025-TP53 library as an example, shown here is a fold-change scatter plot comparing fold-change calculated by raw counts versus that by scaled data. The concordance between the two are apparent.

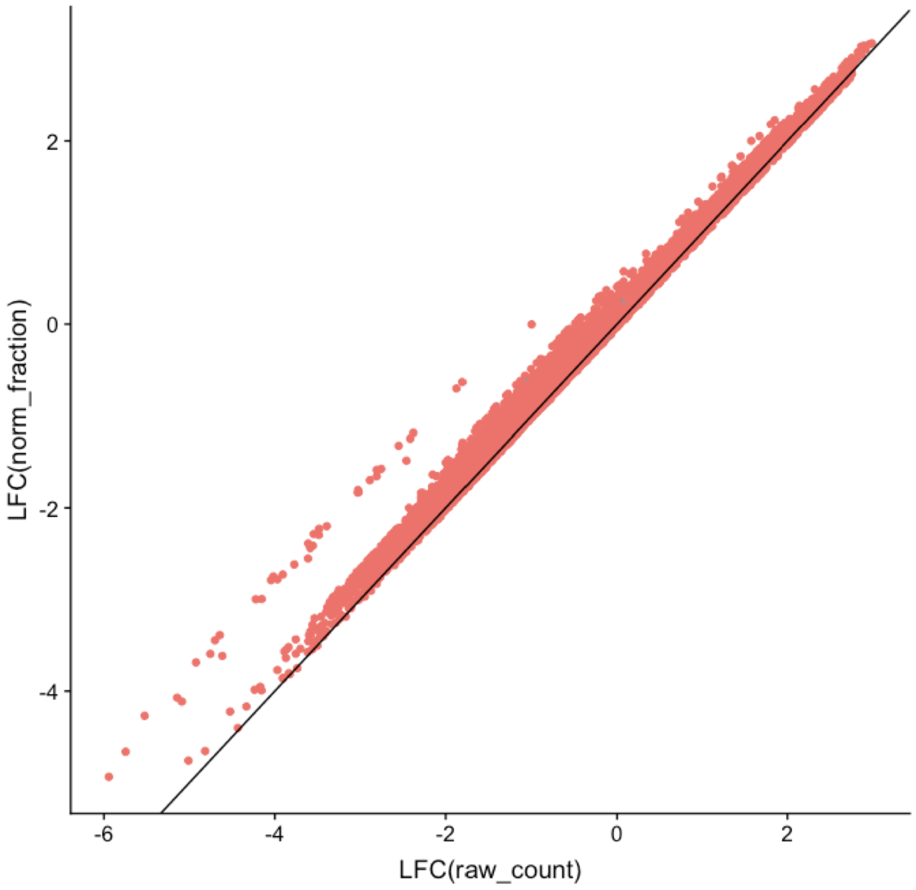
